# Organelle Aging Predicts Human Disease

**DOI:** 10.64898/2026.11.01.763400

**Authors:** Kejun Ying, Kyle KunHyung Roh, Kenneth L Chen, Veronica Augustina Bot, Daisy Yi Ding, Amelia Farinas, The Global Neurodegeneration Proteomics Consortium (GNPC), Carl Ma, Hamilton Se-Hwee Oh, Alina Isakova, Tony Wyss-Coray

## Abstract

Aging affects cellular organelles differently, but whether subcellular aging predicts human disease remains unknown. Here, we integrated organelle-enriched proteins in plasma and cerebrospinal fluid proteomes from 61,931 individuals to model aging of 10 organelles across the human lifespan. Organelle aging was heterogeneous with lysosomes, plasma membranes, and Golgi compartments accelerating rapidly after age 60. Individuals with four or more prematurely aged compartments exhibited up to 22-fold higher kidney disease risk and up to 8-fold higher mortality risk. Major diseases are characterized by distinct organelle aging signatures and disease risk increased with the number of accelerated organelles. APOE-ε4 dosage was associated with lysosomal and mitochondrial aging and reduced PDP1 levels, implicating altered energy metabolism. Decomposition of organelle signature coefficients into biological programs further showed that diseases are associated with distinct subcellular patterns. These findings establish organelle-resolved aging as a measurable dimension of human biology, revealing subcellular mechanisms underlying disease susceptibility and resilience.

One sentence summary: Human organelles age at different rates, revealing distinct molecular signatures that predict disease risk, resilience and mortality.

## Introduction

Aging is the primary risk factor for most chronic diseases, yet the specific molecular processes that drive age-related decline remain incompletely understood. Previous studies have shown that regression models trained on epigenetic (*1–4*) or proteomic (*5–9*) data can estimate biological age and that deviations from chronological age predict mortality and disease risk independently of calendar age (*10–13*). Recent work has extended this approach beyond whole-organism aging to organ- and cellular-level resolution. Proteins enriched in specific organs and detectable in plasma carry distinct aging signatures, and organ-specific age acceleration predicts corresponding organ diseases (*14–16*). Ding et al. further demonstrated that cell-type-specific aging signatures can be extracted from plasma proteomics and predict disease and mortality with cellular specificity (*17*). Such hierarchy of aging phenotypes spanning organisms, organs and cell types raised the question whether aging also varies within cells, across subcellular compartments, where molecular processes maintaining cellular homeostasis are spatially organized.

Strong biological evidence exists to expect such heterogeneity, as the hallmarks of aging include mitochondrial dysfunction, loss of proteostasis, disabled macroautophagy, and deregulated nutrient sensing (*18*), all processes maintained by organelles. Indeed, mutations disrupting organelle functions directly cause numerous human diseases (Table 1), and organelle dysfunctions are linked with aging. Lysosomal dysfunction has been linked to neurodegeneration and metabolic disease (*19, 20*), endoplasmic reticulum stress contributes to insulin resistance and hepatic injury (*21, 22*), and plasma membrane remodeling accompanies cellular senescence (*23, 24*). Autophagy, coordinated by the lysosomal master regulator TFEB (*25, 26*), declines with age across tissues (*27*). Yet despite compelling evidence that organelle dysfunction drives aging and disease, we lack a systematic understanding of how individual organelles age in humans, whether they age independently, and how their distinct aging trajectories contribute to disease risk.

**Table 1:** Mutations in organellar genes directly cause human disease. Examples of disease-associated genes, their subcellular compartments and the principal biological processes or phenotypes affected. Abbreviations: SMA-PME, spinal muscular atrophy with progressive myoclonic epilepsy; VLCFA, very-long-chain fatty acid; CMT, Charcot–Marie–Tooth disease; CDG, congenital disorder of glycosylation; SOPH, short stature, optic atrophy and Pelger–Huët anomaly; BPAN, β-propeller protein-associated neurodegeneration; NDD, neurodevelopmental disorder; AD, autosomal dominant; ER, endoplasmic reticulum; COPII, coat protein complex II; SNARE, soluble N-ethylmaleimide-sensitive factor attachment protein receptor.

| Organelle | Gene examples | Genetic disease | Dominant biology / phenotype |
|---|---|---|---|
| Lysosome | GBA1 | Gaucher disease *(59)* | Glucosylceramide degradation; visceral disease w/ neurodegeneration |
|  | ASAH1 | Farber disease; SMA-PME | Ceramide degradation; inflammation, motor neuron disease, epilepsy |
|  | RNASET2 | RNASET2-deficient cystic leukoencephalopathy *(60)* | Lysosomal RNA degradation; white-matter disease, neurodegeneration |
|  | GLB1 | GM1 gangliosidosis | Ganglioside degradation; progressive neurodegeneration |
|  | HEXA | Tay-Sachs disease | GM2 degradation; severe neurodegeneration |
|  | NPC1 / NPC2 | Niemann-Pick type C | Lysosomal cholesterol export; ataxia, dementia, neurodegeneration |
|  | MCOLN1 | Mucolipidosis IV | Lysosomal ion channel/trafficking; severe neurodevelopmental disease |
|  | LAMP2 | Danon disease | Lysosome/autophagy; cardiomyopathy, myopathy |
|  | CTNS | Cystinosis | Lysosomal cystine export; kidney/eye disease |
| Peroxisome | PEX1, PEX6, PEX12, etc. | Zellweger spectrum disorders | Peroxisome biogenesis; brain, liver, eye, hearing abnormalities |
|  | ABCD1 | X-linked adrenoleukodystrophy | VLCFA transport; inflammatory cerebral demyelination/adrenal disease |
|  | PHYH / PEX7 | Refsum disease | Phytanic-acid α-oxidation; neuropathy, ataxia, retinitis pigmentosa |
| Mitochondria | POLG | POLG-related disorders | mtDNA replication; epilepsy, ataxia, neuropathy |
|  | OPA1 | Dominant optic atrophy | Mitochondrial fusion/cristae; retinal ganglion-cell degeneration |
|  | MFN2 | CMT2A *(61)* | Mitochondrial fusion/transport; peripheral axonal degeneration |
|  | PINK1 / PRKN | Familial Parkinson disease | Mitochondrial quality control/mitophagy |
|  | TWNK | Progressive external ophthalmoplegia | mtDNA maintenance |
| Golgi | COG1/4/5/6/7/8 | COG-CDGs | Golgi trafficking/glycosylation; multisystem developmental disease |
|  | TMEM165 | TMEM165-CDG | Golgi ion homeostasis/glycosylation; skeletal + neurological disease |
|  | ATP6V0A2 | Cutis laxa type II / CDG | Golgi acidification and trafficking |
|  | GOSR2 | Progressive myoclonus epilepsy | Golgi SNARE trafficking; epilepsy/ataxia |
|  | RAB33B | Smith-McCort syndrome | Golgi trafficking; skeletal dysplasia |
|  | GORAB | Gerodermia osteodysplastica | Golgi trafficking; premature-appearing skin/osteoporosis |
| ER / ER–Golgi | SEC23A | Cranio-lenticulo-sutural dysplasia *(62)* | COPII ER-Golgi transport |
|  | SAR1B | Chylomicron retention disease | ER-Golgi lipid transport |
|  | CLN6 / CLN8 | Neuronal ceroid lipofuscinosis | ER export of lysosomal enzymes; progressive neurodegeneration |
|  | NBAS | SOPH syndrome / recurrent acute liver failure | ER-Golgi retrograde trafficking |
| Endosome | VPS35 | Familial Parkinson disease *(63)* | Retromer/endosomal sorting |
|  | RAB7A | CMT2B | Late-endosomal trafficking; peripheral neuropathy |
|  | WASHC5 | Ritscher-Schinzel/3C-like syndrome | Endosomal actin/retromer trafficking |
| Autophagosome | WDR45 | BPAN | Autophagosome formation; childhood NDD followed by adult neurodegeneration |
|  | ATG5 / ATG7 | Neurodevelopmental syndromes | Autophagy; developmental delay, ataxia |
| Nucleus / nuclear envelope | LMNA | Progeria, muscular dystrophies, cardiomyopathy *(64)* | Nuclear lamina architecture |
|  | EMD | Emery-Dreifuss muscular dystrophy | Nuclear envelope |
| Cilium | BBS genes | Bardet-Biedl syndrome | Primary-cilium trafficking/signaling |
|  | IFT140 | Mainzer-Saldino syndrome | Intraflagellar transport |
|  | PKD1 / PKD2 | AD polycystic kidney disease | Ciliary signaling/ion transport |

Herein, we developed organelle-resolved aging models using plasma proteomics across three cohorts. We assigned proteins to 10 subcellular compartments using graph-based organelle annotation applied to native immunoprecipitation proteomics (*28*) and trained separate models to predict age and mortality risk for each organelle, in the UK Biobank (*29*) and the Global Neurodegeneration Proteomics Consortium (GNPC) (*30*) plasma and CSF cohorts (*31*). We show that organelle age acceleration predicts disease risk with organelle-specific patterns across cohorts and biofluids, and that unique patterns of extreme multi-organelle aging confer dramatically elevated risk for specific diseases. Finally, we used differences in aging velocity and the decomposition of organelle signature coefficients into functional modules to reveal biologically interpretable programs linking subcellular aging signatures to human diseases.

## Results

### Organelle models capture age and mortality-associated variation

We analyzed plasma and cerebrospinal fluid (CSF) proteomic data from three cohorts: UK Biobank (UKB, n = 41,209, ages 40-70) (*29*), GNPC Plasma (n = 19,051, ages 20-90), and GNPC CSF (n = 1,671, ages 30-90), all with balanced sex distributions (Fig. 1B). Proteins were assigned to subcellular compartments using graph-based annotations from Hein et al., who used native organelle immunoprecipitation coupled to mass spectrometry and k-nearest-neighbor graph clustering to localize over 7,600 proteins across 19 compartments (*28*). UKB plasma was measured on the Olink Explore 3072 antibody-based platform (*32*), whereas plasma and CSF in GNPC were measured with the SomaScan v4.1 7K aptamer-based assay. The full annotation encompasses 10 major compartments used in our study: plasma membrane (n = 626 proteins), cytosol (n = 1,759), nucleus (n = 1,665), ER (n = 722), ERGIC (n = 54), Golgi (n = 290), lysosome (n = 165), mitochondrion (n = 838), endosome (n = 305), and cytoskeleton (n = 304) (Fig. 1A). The subset of annotated proteins detectable in the Olink panel was used for model training, ranging from 4 (ERGIC) to 169 (cytosol) proteins per organelle in UKB. Organelle-assigned proteins show significant within-compartment connectivity enrichment (1.2- to 28.4-fold over random) based on protein-protein interaction analysis (*33*) (See Materials and Methods), supporting the biological coherence of these assignments (Fig. 1C).

**Fig. 1:**
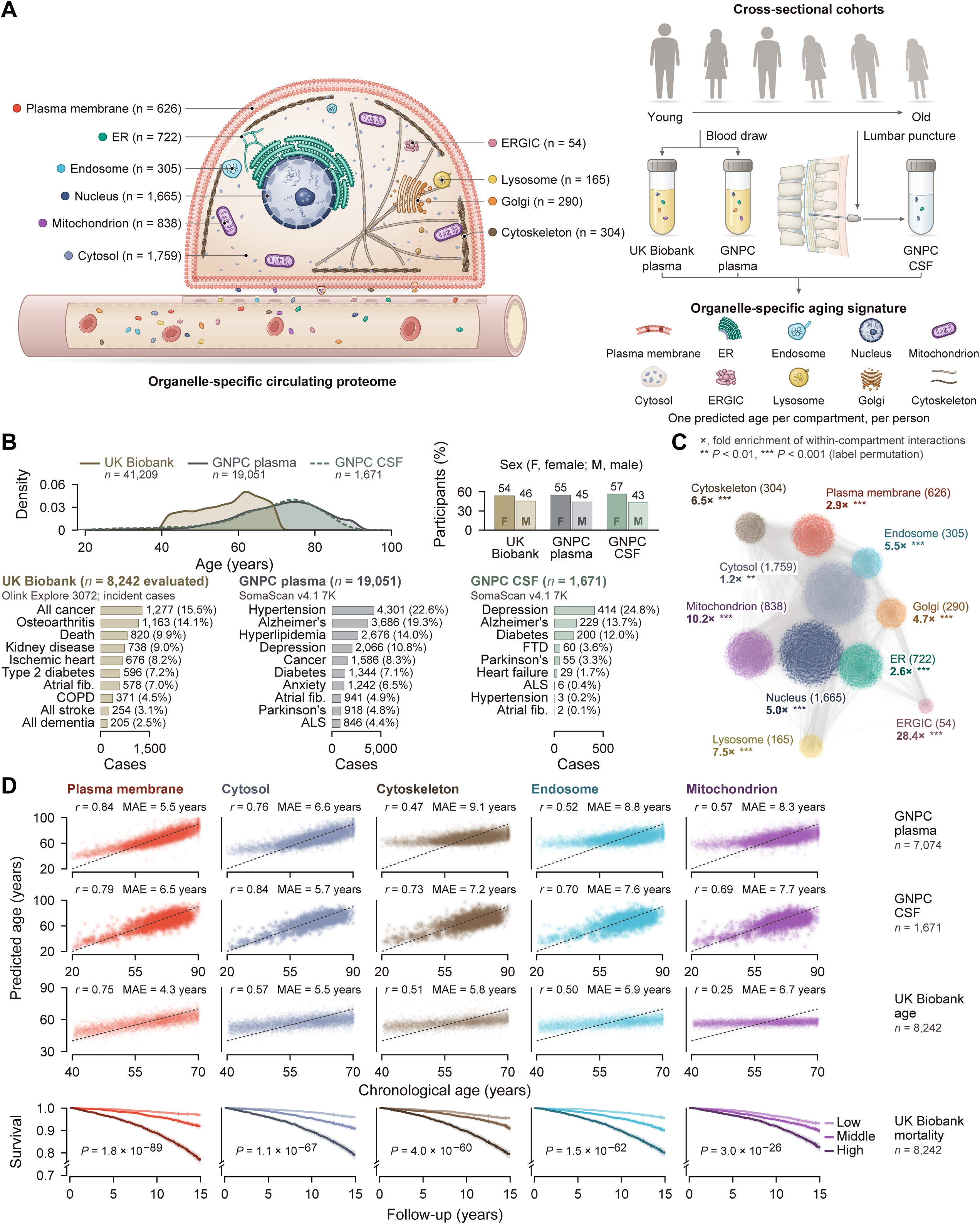
Organelle-resolved proteomics and organelle aging models. (A) Subcellular protein assignments and the strategy for deriving organelle-associated aging signatures from biofluid proteomes. Numbers denote annotated proteins. (B) Age distributions, sex distributions and disease counts in UK Biobank (UKB; n = 41,209), Global Neurodegeneration Proteomics Consortium (GNPC) plasma (n = 19,051) and GNPC cerebrospinal fluid (CSF; n = 1,671). (C) Protein-protein interaction network colored by compartment, with within-compartment connectivity enrichment assessed using STRING (*33*). (D) Predicted versus chronological age for five compartments in GNPC plasma, GNPC CSF and the UKB test set, followed by UKB Kaplan–Meier survival curves stratified by mortality-model risk tertile. Pearson correlation coefficients and mean absolute errors are displayed for age prediction; log-rank P values are displayed for survival comparisons. Models were trained separately in each cohort. ER, endoplasmic reticulum; ERGIC, ER–Golgi intermediate compartment.

Over half of the measured proteins showed significant age associations (818 of 1,136 in UKB, 674 of 793 in GNPC CSF and 5,567 of 6,219 in GNPC plasma, at p < 0.05; fig. S1, A to C). We next used a two-stage modeling framework to determine whether organelle-associated proteomic signatures capture both chronological aging and mortality risk. First, we trained elastic-net models to predict chronological age from proteins assigned to each organelle. Models were developed using an 80/20 training-test split in UKB. Predictive performance varied substantially across organelles: the plasma-membrane model achieved the lowest mean absolute error (MAE = 4.34 years; r = 0.75), followed by the cytosolic (MAE = 5.53 years; r = 0.57) and cytoskeletal models (MAE = 5.80 years; r = 0.51). By contrast, mitochondrial and nuclear models showed more modest performance (r = 0.25 and 0.21, respectively), potentially due to limited detection of mitochondrial proteins in plasma (Fig. 1D). Models trained independently in GNPC plasma reproduced this relative performance, with the plasma-membrane signature again providing the most accurate age estimates (r = 0.84). Organelle-associated age signatures were also detectable in GNPC cerebrospinal fluid, where plasma-membrane and cytosolic models achieved correlations of r = 0.79 and 0.84, respectively (Fig. 1D).

We then asked whether deviations from these proteomic age estimates were associated with mortality. For each organelle, age acceleration was calculated as the residual from the corresponding predicted-versus-chronological-age relationship and evaluated using L1-penalized Cox regression in UKB. All organelle-specific models achieved C-indices above 0.70, with the strongest discrimination observed for cytoskeletal (C = 0.727), lysosomal (C = 0.723) and endosomal (C = 0.722) signatures. Consistently, Kaplan–Meier (KM) curves stratified by predicted-risk tertile showed significant survival differences across the plasma-membrane, cytosolic, cytoskeletal, endosomal and mitochondrial models (all log-rank P < 0.001; Fig. 1D). Thus, deviations in compartment-specific proteomic age were associated with subsequent mortality, establishing the clinical relevance of these signatures.

Having established their prognostic relevance, we next tested whether organelle-specific age gaps reflected a common systemic aging process or distinct compartmental programs. Pairwise correlation analysis revealed a combination of shared and compartment-selective variation (r = 0.2–0.6). In GNPC plasma, components of the secretory pathway, including the endoplasmic reticulum, ER–Golgi intermediate compartment and Golgi apparatus, were more strongly correlated with one another than with degradative compartments such as lysosomes and endosomes, but this pattern was absent in UKB and GNPC CSF. Correlations were also stronger overall in GNPC plasma than in UKB (fig. S2A). Organelle aging is therefore only partly shared, and a substantial fraction of each organelle’s aging signal is independent of the others. At the protein level, features selected by the age-prediction models showed similarly structured changes, including coordinated age-associated increases among endosomal, lysosomal, Golgi and ERGIC proteins and decreases among mitochondrial and nuclear proteins (fig. S2B). These analyses established that organelle-resolved proteomic signatures generalize across independent cohorts and biofluids, and therefore we used the computed age gaps to test whether human disease maps to distinct subcellular aging states.

### Disease associations vary across organelles and biofluids

Organelle age acceleration in GNPC plasma showed significant compartment-specific risk across neurodegenerative diseases, cardiometabolic, cancer, and COPD. For example, Alzheimer’s disease (AD) was significantly associated with Golgi, plasma membrane, and mitochondrion, while frontotemporal dementia (FTD) was strongly associated with lysosome (Fig. 2A). Given the strong genetic links between FTD and lysosomal dysfunction, including mutations in *GRN, CHMP2B*, and *C9orf72*, as well as risk variants in *TMEM106B* (*34*), our findings demonstrate the ability of organelle-resolved aging signatures to identify disease-relevant biological pathways. Individuals with extremely accelerated or decelerated organelle aging exhibited significantly increased or decreased disease risk, respectively, and accelerated aging was associated with higher mortality (Fig. 2, B and C). Notably, extreme plasma membrane agers had markedly elevated risk for kidney disease (HR = 2.77, p = 3.5 x 10^−10^; fig. S3, A to C).

**Fig. 2:**
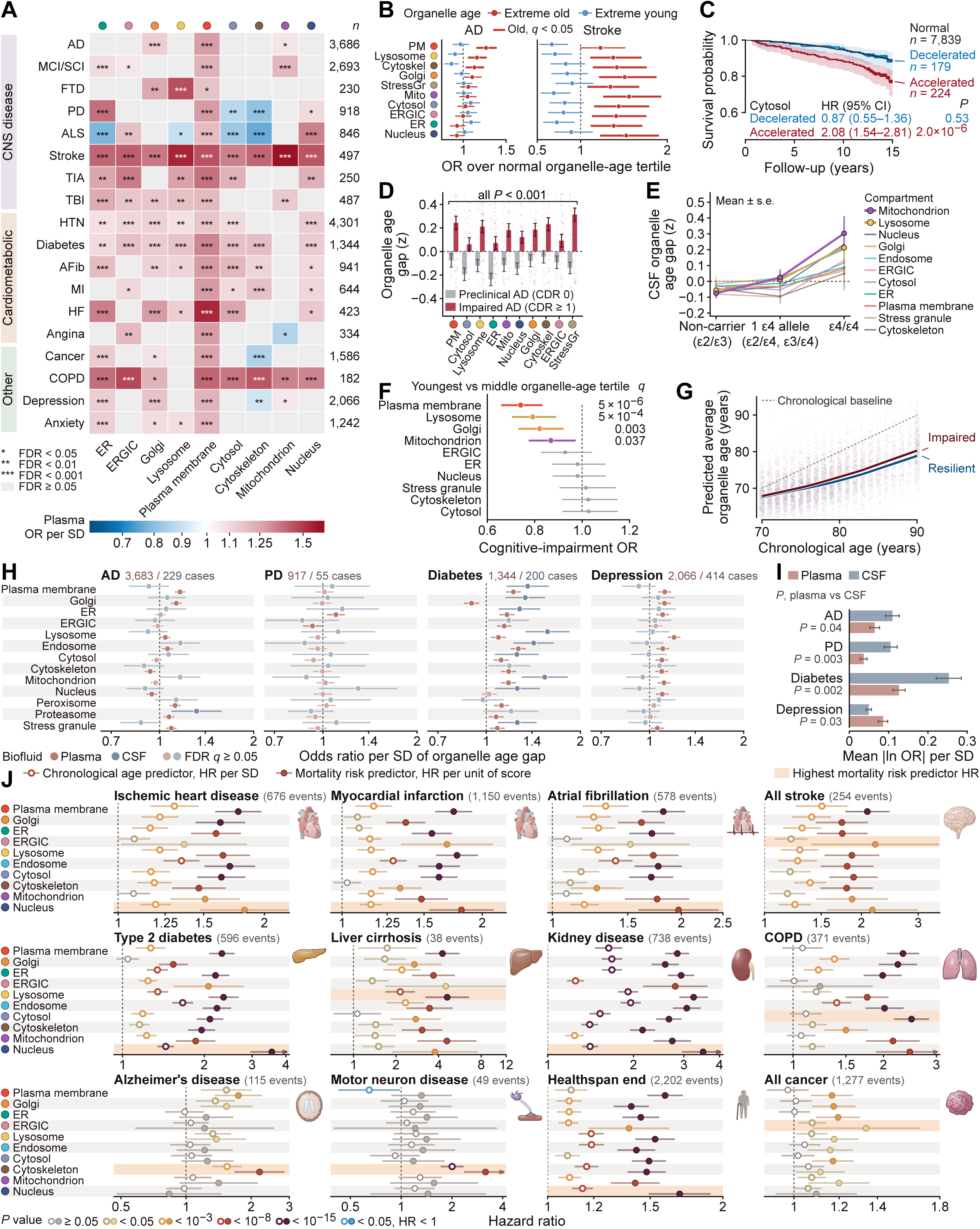
Organelle age acceleration is associated with disease and cognitive status. (A) Organelle age-gap associations with prevalent diseases in GNPC plasma, where colors indicate odds ratios (ORs) per SD of age gap, gray cells have FDR ≥ 0.05, and asterisks indicate false discovery rate thresholds shown below the heatmap. (B) ORs for Alzheimer’s disease (AD) and stroke comparing older and younger organelle-age tertiles with the middle tertile. (C) UKB survival by cytosolic aging group (accelerated, age gap > 2 SD; decelerated, < −2 SD) in the held-out test set, with hazard ratios (HRs) from Cox models adjusted for age and sex. (D) Mean organelle age acceleration and 95% confidence intervals (CIs) in recorded AD cases with Clinical Dementia Rating (CDR) 0 or ≥1; all compartments differ between the groups (two-sided Mann-Whitney P < 0.001). (E) CSF organelle age gaps by number of APOE-ε4 alleles, points and bars show means and standard errors. (F) Age- and sex-adjusted cognitive-impairment ORs for the youngest versus middle organelle-age tertile, with 95% CIs and adjusted q values. (G) Predicted organelle age versus chronological age in impaired and resilient groups. (H) Plasma and CSF ORs per SD of organelle age gap and 95% CIs for four diagnoses, from logistic regression adjusted for age and sex; faded points have FDR ≥ 0.05 (Benjamini-Hochberg across the 52 tests in each biofluid), and numbers give cases in plasma and CSF. (I) Mean absolute log(OR) across the 13 compartments by biofluid; error bars show s.e.m., and P values are from two-sided paired t-tests across compartments. (J) UKB incident-disease HRs and 95% CIs for chronological-age predictors (per SD) and mortality-risk predictors (per unit of score); shading marks the compartment with the highest mortality-predictor HR.

Organelle age differed across stages of AD, where the number of APOE-ε4 alleles significantly correlated with acceleration in mitochondria and lysosomes (Fig. 2, D and E), aligning with prior experimental observations in the field that link APOE-ε4 to impaired lysosomal function and mitochondrial bioenergetics (*35–37*). Conversely, deceleration in organelle ages in plasma membrane, lysosome, Golgi, and mitochondria was strongly associated with cognitive resilience, and this pattern was consistent from 70 to 90 years of age (Fig. 2, F and G).

Indeed, APOE-ε4 may influence lipid availability and immune regulation across lifespan, raising the possibility that its association with mitochondrial aging partly reflects an underlying immunometabolic phenotype (*38–42*). To distinguish the genotype-associated proteomic variation from that accompanying AD, we decomposed the mitochondrial signature and examined its contributors across disease status. PDP1 made the largest individual contribution to the AD-adjusted association between ε4 dosage and mitochondrial age acceleration (fig. S4A). More ε4 alleles were associated with lower CSF PDP1 abundance despite conditioning on age, sex, recorded AD status, 36 immune/lipid proteins and four APOE assays (β = −0.410 SD per allele, 95% CI −0.507 to −0.313; q = 1.32 × 10⁻¹⁴; fig. S4B).

The association was present in participants both without and with recorded AD (β = −0.409 and −0.442, respectively), with no evidence of modification by AD status (interaction P = 0.915; fig. S4, C and D), suggesting that mitochondrial protein signatures sufficiently distinguish effects of APOE-ε4 beyond association with AD and immune response. APOE-ε4 was also associated with modestly accelerated lysosomal aging, largely driven by the lysosomal protein ASAH1 (fig. S5). Among ε4 carriers, higher levels of CREG1, BNIP3L and SIRT5 were associated with lower AD risk, but these associations were no longer significant after correcting for multiple comparisons (fig. S6).

Disease associations in GNPC were concordant with those observed in UKB. Notably, organelle aging signatures in CSF showed particularly strong associations with neurological diseases, including AD and Parkinson’s disease (PD) (*31, 43*), whereas plasma-derived signatures were associated with a broader range of diseases, highlighting the distinct information captured by each biofluid (Fig. 2, H to J). Interestingly, across all diseases examined, diabetes showed the strongest risk association with accelerated aging of an individual organelle.

To identify molecular pathways linking organelle aging with disease, we examined which specific proteins contributed most strongly to organelle aging signatures associated with specific diseases. Proteins with both large aging signature coefficients and strong associations with disease risk were considered candidate mechanistic links. In AD, cytoskeletal proteins including NEFL, a well-established biomarker of neuroaxonal damage (*44*), BRK1, and NCAM2 contributed prominently and pointed to enrichment in pathways including “regulation of actin cytoskeleton” (p = 2.0 x 10^−4^). In liver cirrhosis, lysosomal proteins PLA2G15, CTSL, and SCARB2 were linked to apoptosis pathways (p = 1.2 x 10^−5^). In COPD, cytosolic proteins CSTB, LGALS1, and GLO1 were linked to tuberculosis-related pathways, whereas healthspan endpoints were linked to nuclear proteins DPY30 and ZBTB17, which are involved in transcriptional regulation and enriched in cancer pathways (p = 7.0 x 10^−3^; fig. S3, D to G).

### Multi-organelle aging and discordance are associated with disease

Because aging may affect multiple organelles within an individual, we investigated whether the extent and pattern of subcellular age acceleration can provide additional information about disease risk and clinical significance. t-SNE visualization revealed distinct groups of individuals characterized by accelerated aging of unique organelles including the plasma membrane, mitochondrion, nucleus, lysosome, and ER, as well as individuals with accelerated aging across multiple organelles (Fig. 3, A and B). Strikingly, disease risk increased with the number of extremely aged organelles (Fig. 3C). Individuals with accelerated aging of a single organelle had modestly elevated risks (HR = 1.3–2.0 for most outcomes), while those with four or more prematurely aged organelles showed up to 22-fold higher kidney disease risk and up to 8-fold higher mortality risk. Similar associations were observed in GNPC plasma for 12 of 18 prevalent diseases and in GNPC CSF for frontotemporal dementia, heart failure and diabetes (Fig. 3D).

**Fig. 3:**
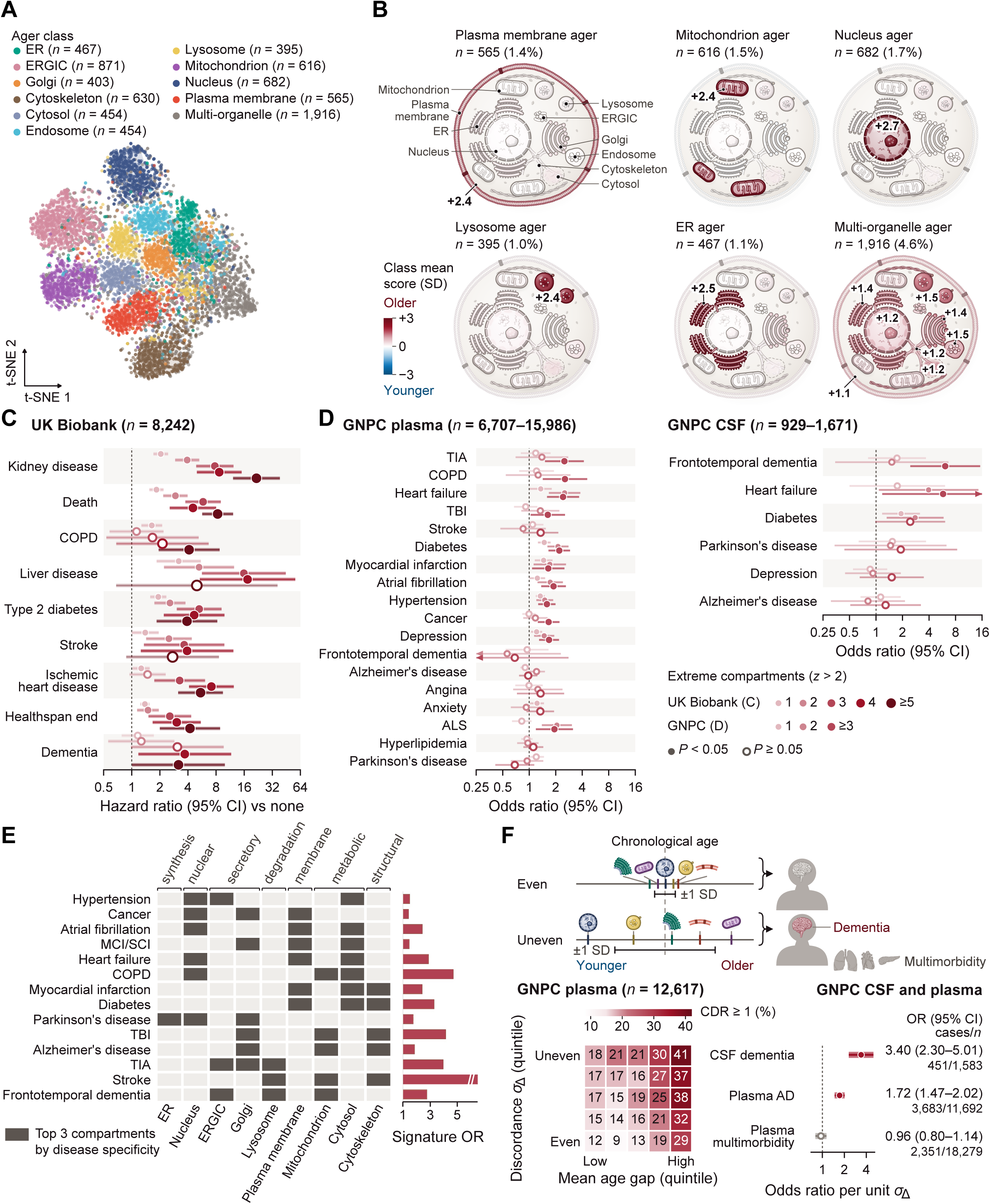
Multi-organelle age acceleration and discordance are associated with disease. (A) t-distributed stochastic neighbor embedding of participants with extreme organelle age acceleration, colored by aging profile. (B) Compartment-specific mean age-gap profiles for six illustrated ager groups; sample counts and proportions are shown. (C) UKB disease HRs by the number of simultaneously extreme organelles (z > 2), relative to participants with none, in the held-out test set with adjustment for age and sex. (D) Corresponding prevalent-disease ORs in GNPC plasma and CSF. Horizontal intervals in C and D indicate 95% CIs. (E) Disease-associated organelle signatures, with diseases in hierarchical-clustering order. Dark cells mark the three compartments with the highest disease specificity, column labels give functional categories, and bars give the associated signature ORs. (F) Illustration of within-person organelle discordance, the prevalence of dementia across joint bins of mean age gap and dispersion, and discordance-associated ORs with 95% Wald CIs. Discordance is the within-person SD across standardized organelle age gaps. Logistic models include mean age gap, linear and quadratic age terms, and sex. AD, Alzheimer’s disease; CSF, cerebrospinal fluid.

We next asked if specific combinations of aged organelles were associated with different diseases. Hierarchical clustering revealed distinct multi-organelle aging signatures (Fig. 3E). Cardiovascular and cerebrovascular disorders shared broad aging signatures involving membrane, secretory, nuclear, and degradative compartments. Neurodegenerative diseases were more strongly associated with secretory, degradative, metabolic, and cytoskeletal compartments.

Finally, we asked whether discordance in aging among organelles within the same individual was associated with disease, independent of overall age acceleration. Greater discordance between organelle aging, with some compartments appearing biologically older and others younger, was associated with dementia in CSF and with AD in plasma (Fig. 3F). This association was not observed for multimorbidity, suggesting that disease risk depends not only on the extent of organelle aging but also on the specific configuration of aging across subcellular compartments.

### Aging velocities differ across subcellular compartments

Having established that organelle aging estimates predict disease and mortality, we next investigated whether the proteins underlying organelle signatures could provide mechanistic insights into aging. First, we combined the proteins assigned to each organelle into a single standardized aging signature and examined how these signatures changed across the adult lifespan (Fig. 4A). Although all nine compartment signatures showed similar trajectories through early midlife, their aging patterns diverged between approximately 57 and 63 years of age. Notably, the lysosome, plasma membrane, and Golgi apparatus (LPG) exhibited substantially faster aging than the remaining six compartments (Fig. 4B). We quantified aging velocity as the change in each standardized signature per decade, with the lysosomal signature increasing by roughly one-third of a standard deviation per decade (Fig. 4B). The mean aging velocity of these three LPG compartments exceeded that of the other six compartments in every bootstrap resample (p < 0.001, Fig. 4B).

**Fig. 4:**
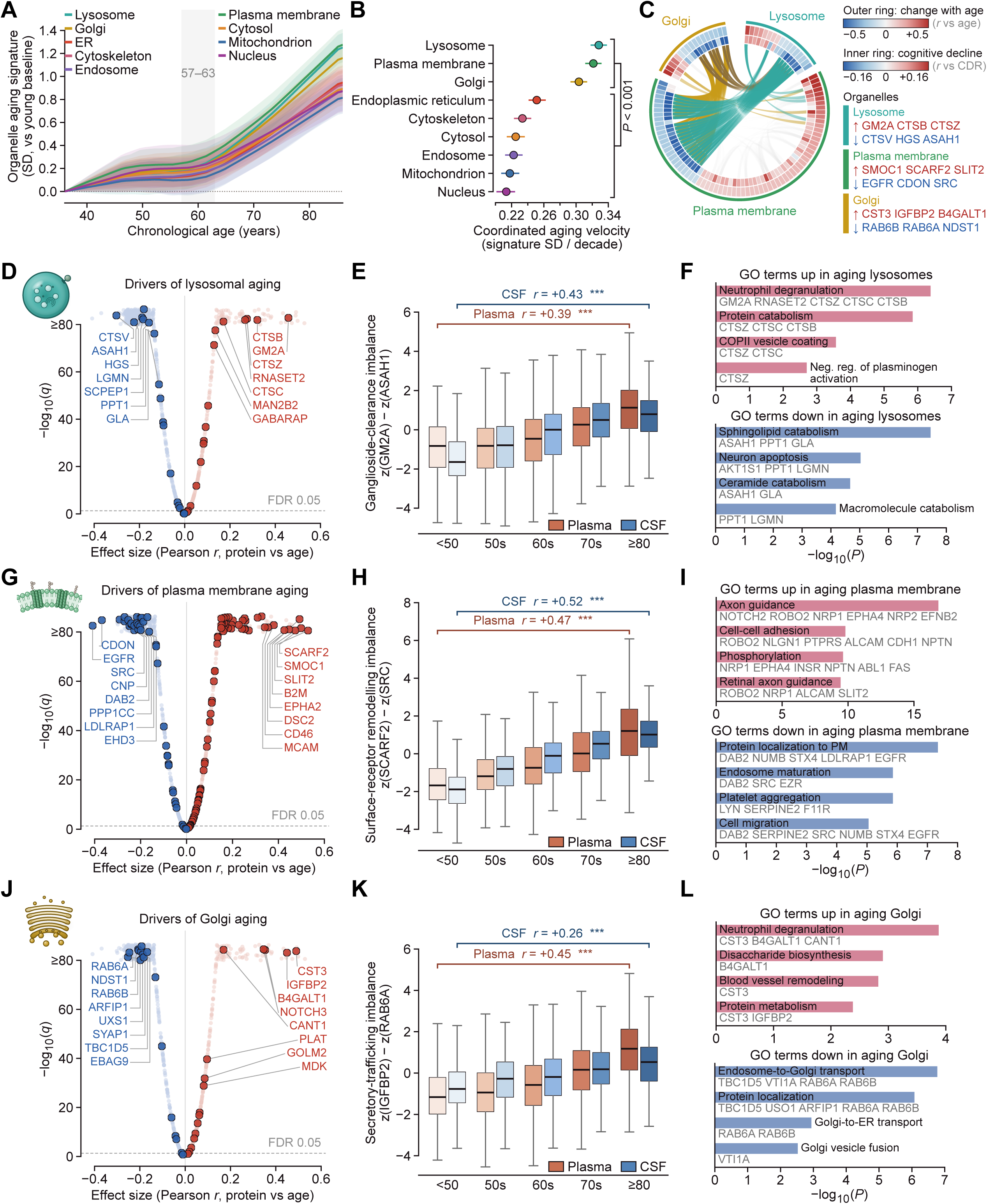
Differential aging velocity identifies coordinated lysosomal, plasma-membrane and Golgi signatures. (A) Smoothed organelle protein signatures across age in GNPC plasma, referenced to the young baseline; shading shows bootstrap 95% confidence bands. (B) Signature change per decade and subject-bootstrap 95% CIs, with the lysosome–plasma membrane–Golgi (LPG) group compared with the other six compartments. (C) Cross-compartment protein correlations after age detrending. Outer and inner tracks show age correlations and age-controlled correlations with CDR, respectively. (D) Lysosomal protein–age correlations and −log10(P), with the false discovery rate threshold indicated. (E) Ganglioside-clearance imbalance, z(GM2A) − z(ASAH1), across age groups in plasma and CSF. (F) Enriched Gene Ontology terms among age-increasing and age-decreasing lysosomal proteins. (G– I) Corresponding plasma-membrane analyses, including the imbalance z(SCARF2) − z(SRC). (J–L) Corresponding Golgi analyses, including z(IGFBP2) − z(RAB6A). In E, H and K, boxes show medians and interquartile ranges, and whiskers extend to the most extreme observations within 1.5 interquartile ranges. Pearson r describes association with age; asterisks denote two-sided Mann–Whitney comparisons of the youngest and oldest bins (***P < 0.001). Pathway bars show −log10(P).

We then asked whether these three rapidly aging LPG compartments were part of a coordinated program. Analysis of age-adjusted protein correlations revealed coordinated changes in LPG-associated proteins that were also linked with cognitive decline (Fig. 4C). Consistent with this observation, the three LPG protein signatures remained strongly correlated after nonlinear age adjustment in 19,051 GNPC plasma participants, with residual correlations of 0.81 between lysosome and plasma membrane, 0.79 between lysosome and Golgi, and 0.89 between plasma membrane and Golgi (fig. S7). Five-protein signatures selected by their age correlations from each compartment showed distinct associations with neurological and cerebrovascular diseases (fig. S8), linking this coordinated aging program to clinical phenotypes. Thus, we identified the main molecular drivers within each of the LPG compartments (Fig. 4D, G, J).

Within lysosomes, aging was associated with significantly increased levels of cathepsins and the GM2 ganglioside activator GM2A, while the sphingolipid hydrolases ASAH1, PPT1, and GLA decreased (Fig. 4D). These opposing changes suggest an age-related imbalance in glycosphingolipid processing. To quantify this imbalance, we calculated the difference between standardized GM2A and acid ceramidase ASAH1 levels, which increased monotonically with age in plasma (r = 0.39) and CSF (r = 0.43) (Fig. 4E). In agreement with these findings, gene ontology (GO) analysis revealed enrichment of neutrophil degranulation and protein catabolism among proteins increasing with age and sphingolipid and ceramide catabolism among proteins decreasing with age (Fig. 4F).

We observed similar patterns of divergence between increasing and decreasing proteins in plasma membrane and Golgi aging (Fig. 4G, J). At the plasma membrane, scavenger and adhesion receptors, including SCARF2, SMOC1, and MCAM, increased with age, while the signaling components EGFR, SRC, and DAB2 decreased (Fig. 4G), pointing to upregulation of axon guidance and cell-to-cell adhesion and downregulation of protein localization to the plasma membrane and endosome maturation with age (Fig. 4I). A synthetic surface-receptor-remodeling index defined as the difference between SCARF2 and SRC levels increased steadily with age and showed strong correlations in both plasma (r = 0.47) and CSF (r = 0.52), suggesting brain and periphery might be similarly susceptible to age-related plasma membrane deterioration (Fig. 4H). At the Golgi apparatus, secreted cargo and glycosylation enzymes including IGFBP2, CST3, and B4GALT1 increased and Rab GTPases RAB6A and RAB6B, involved in retrograde trafficking, decreased with age (Fig. 4J) and pointed to enrichment in secretory and degranulation programs and downregulation of endosome-to-Golgi and Golgi-to-endoplasmic-reticulum transport (Fig. 4L). An index capturing this secretory-trafficking imbalance defined as the difference between IGFBP2 and RAB6A levels increased with age as well (Fig. 4K). Taken together, these findings suggest a coordinated aging signature across the lysosome, plasma membrane, and Golgi apparatus, characterized by declining lipid-processing and intracellular trafficking proteins alongside increasing inflammatory, secretory, and degranulation-associated proteins. All three molecular imbalance indices showed concordant age-related changes in plasma and CSF (Fig. 4E, H, and K). The observations are consistent with experimental findings that age-related decline in lysosomal degradation can lead to the accumulation of cholesterol and sphingolipids, potentially disrupting plasma membrane homeostasis and Golgi trafficking (*23, 27, 45–48*).

### Functional modules organize organelle-associated disease risk

To investigate how organelle aging contributes to disease risk, we first used principal component analysis (PCA) to examine the organization of compartment age gaps in UKB (Fig. 5A). The first principal component (PC1), explaining 36% of the variance, loaded positively on every compartment and was dominated by the plasma membrane. PC2, explaining an additional 16%, separated the plasma membrane from all other compartments. Because these components reflect compartment variance as much as function, we grouped compartments into functional modules by their known biology.

**Fig. 5:**
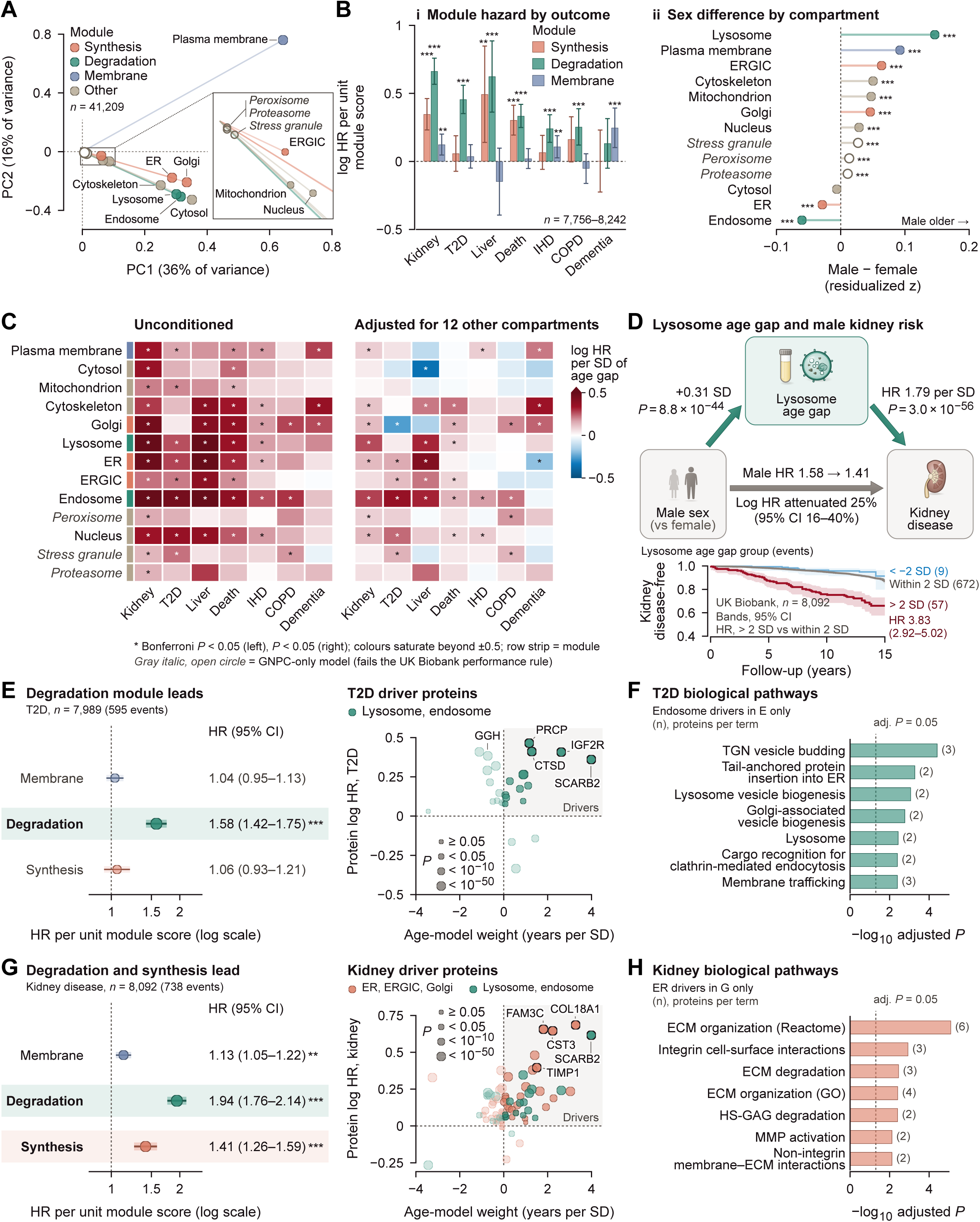
Functional organization of organelle signatures and disease-associated protein programs. (A) Principal component analysis of UKB compartment age gaps (n = 41,209); points show compartment loadings on PC1 and PC2, colored by functional module. (B) Synthesis, degradation and membrane module associations with disease, expressed as log(HR) per unit of module score in the held-out test set, and male–female differences in residualized organelle age gaps at right. (C) Organelle - disease log(HR) estimates before and after adjustment for the 12 other compartment age gaps; asterisks mark Bonferroni-significant associations (left) and P < 0.05 (right), and gray italic labels mark GNPC-only models that did not meet the UKB performance threshold. (D) Statistical accounting of the male-kidney disease association by lysosomal aging, with Kaplan-Meier curves by lysosomal age-gap group (< −2 SD, within 2 SD, > 2 SD). (E) Type 2 diabetes module HRs and 95% CIs per unit of module score, and protein-level disease associations plotted against age-model weights, with point size marking P-value bins. (F) Enriched biological pathways for the endosomal type 2 diabetes driver proteins. (G) Corresponding module and protein analyses for kidney disease. (H) Enriched pathways for the ER kidney disease driver proteins. Protein colors and symbols indicate compartment assignments. Pathway bars show −log10(adjusted P), and parenthetical counts indicate mapped proteins. ER, endoplasmic reticulum; ERGIC, ER-Golgi intermediate compartment.

We defined three functional modules: synthesis (ER, ERGIC, and Golgi), degradation (lysosome and endosome), and membrane (plasma membrane). Cox regression models using these module-level scores revealed distinct patterns of disease risk (Fig. 5B). Type 2 diabetes risk was associated most strongly with the degradation module (aHR = 1.58), with no significant association with the membrane (aHR = 1.04) or synthesis (aHR = 1.06) modules (Fig. 5E). Key contributing proteins included lysosomal enzymes CTSD and PRCP and the endosomal receptor IGF2R, with enrichment for Golgi-associated vesicle and lysosomal pathways (Fig. 5, E and F). Kidney disease showed a broader pattern with significant contributions from all three modules: degradation (aHR = 1.94), synthesis (aHR = 1.41), and membrane (aHR = 1.13) (Fig. 5G). Key contributing proteins spanned multiple compartments including ER (COL18A1, TIMP1), Golgi (CST3, FAM3C), and lysosome (SCARB2), and were enriched for extracellular matrix organization and integrin signaling (Fig. 5, G and H). These findings suggest that different diseases are associated with distinct combinations of subcellular aging programs rather than a uniform acceleration of organelle aging.

We next examined whether individual organelle-disease associations remained significant after accounting for aging in all other compartments (Fig. 5C). Several initially strong associations were attenuated after adjustment, indicating shared information and biology across organelles. In contrast, others, including nucleus-kidney disease and lysosome-liver disease associations, remained robust, suggesting that certain organelles capture disease-relevant information beyond that provided by other compartments.

Finally, we investigated sex differences in organelle aging and their potential contribution to disease risk. Men exhibited significantly accelerated aging of lysosomal, plasma membrane and ERGIC compartments compared with women, while endosome and ER showed the opposite pattern (Fig. 5B). Notably, mediation analysis suggested that lysosomal aging statistically accounts for 25% of the excess kidney disease risk in men (HR decreasing from 1.58 to 1.41 after adjusting for lysosomal age acceleration) (Fig. 5D). Although this analysis does not establish causality, it suggests that sex differences in lysosomal aging may contribute to the higher incidence of kidney disease in men.

### Discussion and Outlook

In this study, we resolved the human aging proteome into subcellular compartments and demonstrated that organelles follow partially independent aging trajectories associated with distinct disease risks. These findings extend biological aging measurements beyond organs and cell types, providing a framework for identifying disease-relevant molecular mechanisms and potentially enabling more targeted approaches to precision medicine. Individuals with accelerated aging across multiple organelles exhibited particularly high disease and mortality risk. Importantly, disease risk was associated not only with the number of prematurely aged organelles, but also with the degree of discordance among their biological ages, suggesting that distinct patterns of subcellular aging capture aspects of systemic dysfunction that are not apparent from global measures of biological age.

Beyond predicting disease risk, our framework provides insights into the molecular organization of aging through analyses of organelle aging velocities and functional programs. We identified the lysosome, the plasma membrane, and the Golgi apparatus as the most rapidly aging compartments, particularly after midlife. Their coordinated remodeling revealed a common pattern characterized by declining biosynthetic programs, intracellular trafficking, and lipid-clearance capacity alongside increasing inflammatory, secretory, and degranulation-associated proteins (Fig. 4). These findings suggest that aging involves coordinated changes across the endolysosomal and secretory systems rather than independent deterioration of individual organelles.

Our approach in modelling organelle aging further revealed associations between APOE-ε4 and mitochondrial and lysosomal aging (Fig. 2E), consistent with previous literature linking APOE4 to impaired organelle function (*35–37*). Notably, analysis of the composition of the mitochondrial organelle signature identified PDP1 as a major contributor to its association with APOE-ε4 (fig. S4), revealing a previously unrecognized link between APOE genotype and this regulator of mitochondrial metabolism. PDP1 activates pyruvate dehydrogenase through dephosphorylation, promoting the conversion of pyruvate to acetyl-CoA and supporting mitochondrial glucose oxidation. Experimental studies have linked this pathway to activity-dependent neuronal energy metabolism (*49, 50*). Consistent with our findings, APOE4-expressing neuronal cells exhibit reduced PDH subunit abundance and impaired respiratory capacity, while isotope-tracing studies in APOE4 models have demonstrated increased glycolytic flux at the expense of glucose oxidation (*37, 51*). The inverse association between APOE-ε4 dosage and CSF PDP1 levels (fig. S4B) therefore suggests that altered regulation of pyruvate oxidation may contribute to the metabolic phenotype associated with APOE4. Similar metabolic reprogramming has been observed in APOE4-expressing microglia, providing a potential link to immune function (*52*). Together, these observations identify PDP1 and the regulation of pyruvate oxidation as potential mechanisms connecting APOE4 to altered cellular metabolism.

Functional decomposition of organelle aging signatures into synthesis, degradation, and membrane modules further uncovered distinct biological programs associated with disease risk. Type 2 diabetes was most strongly associated with accelerated aging of degradative compartments, consistent with established roles of lysosomal dysfunction and impaired autophagy in pancreatic beta-cell failure and insulin resistance (*22, 53*). Kidney disease, on the other hand, was associated with all three modules, suggesting disruptions in protein homeostasis and membrane remodeling. These findings are consistent with reports that lysosomal dysfunction in renal tubular cells promotes senescence and interstitial fibrosis through impaired TFEB-dependent autophagy (*54, 55*). Thus functional decomposition of organelle aging signatures may help identify disease-specific vulnerabilities and generate hypotheses for targeted therapeutic interventions.

We also identified pronounced sex differences in organelle aging, with men showing greater lysosomal, plasma membrane, and ERGIC age acceleration than women. Interestingly, lysosomal aging statistically accounted for approximately 25% of the excess kidney disease risk associated with male sex, suggesting that accelerated lysosomal aging in males may contribute to their higher kidney disease incidence. Lysosomal biogenesis and autophagy are regulated by TFEB (*25*), whose expression is influenced by sex hormone signaling. For example, ablation of androgen and estrogen receptor signaling reduces TFEB levels and impairs autophagosome clearance (*56*), providing a plausible biological connection between sex, lysosomal function, and kidney disease. Whether interventions that enhance lysosomal function could preferentially reduce kidney disease risk in men remains an important question for future studies.

Several limitations should be noted. First, plasma proteomics captures only proteins in circulation and likely underrepresents intracellular compartments such as mitochondria. Consequently, organelle aging signatures reflect changes in circulating proteins associated with specific compartments rather than direct measurements of organelle function. Second, protein localization annotations assign proteins to primary compartments, although many proteins function across multiple organelles, potentially limiting the specificity of individual signatures. Third, our observational associations cannot establish causality, and the mediation analysis identifies statistical relationships rather than demonstrating causal mechanisms (*57*). Fourth, the cohorts were predominantly of European ancestry, and the generalizability of these findings requires validation (*9*). Finally, although genetic studies of centenarians suggest that inherited variation contributes to heterogeneity in aging trajectories (*58*), the genetic determinants of organelle-specific aging remain largely unexplored.

Future experimental studies in cellular and animal models will be essential to determine whether molecular pathways identified here contribute causally to organelle aging and whether they can be therapeutically modified. In particular, interventions targeting glycosphingolipid processing, plasma membrane receptor turnover, or retrograde Golgi trafficking could test whether these processes regulate the corresponding organelle aging signatures. Longitudinal studies with repeated proteomic measurements will also be important for establishing the stability, temporal dynamics, and potential reversibility of organelle-specific aging.

Together, our findings establish organelle-resolved proteomics as a new approach to studying human aging at subcellular resolution. By revealing distinct aging trajectories, coordinated molecular programs, and disease-specific patterns of accelerated organelle aging, this framework moves beyond global measures of biological age toward a more mechanistic understanding of aging and disease. As proteomic technologies and longitudinal cohorts expand, organelle aging signatures may provide a scalable approach to identifying individuals at risk, monitoring biological responses to interventions, and guiding the development of therapies targeting specific subcellular processes. Ultimately, resolving biological aging at the organelle level may help advance precision geroscience toward interventions that preserve cellular function and extend healthy lifespan.

## Acknowledgments

We thank the staff and participants in all studies for their important contributions. We thank A. Poyi-Tsai, P. Moran-Losada and other members of the Wyss-Coray laboratory for feedback and support and H. Zhang and D. Channappa for laboratory management. UKB cohort analyses were conducted using the UKB Resource under Application Number 184058.

## Author contributions

Conceptualization: K.Y., T.W.-C. Methodology: K.Y., K.K.-H.R. Software: K.K.-H.R., K.Y. Formal analysis: K.Y., K.K.-H.R., K.L.C. Investigation: K.Y., K.K.-H.R., V.A.B., D.Y.D., A.F., C.M., H.S.-H.O. Resources: T.W.-C. Data curation: K.K.-H.R., K.L.C., V.A.B. Visualization: K.Y., K.K.-H.R. Supervision: T.W.-C. Funding acquisition: T.W.-C., K.Y. Writing, original draft: K.Y. Writing, review and editing: all authors.

## Data, code, and materials availability

UK Biobank data are available to approved researchers through the UK Biobank Access Management System. GNPC plasma and CSF data are available to approved researchers through the GNPC data access process. Analysis code will be deposited in a public repository before publication. All other data are in the main text or the supplementary materials.

## List of Supplementary Materials

Materials and Methods

Figs. S1 to S8

References (65–74)

Members of the Global Neurodegeneration Proteomics Consortium

