## Supplementary Materials for "Organelle Aging Predicts Human Disease"

‡Members of the GNPC and their affiliations are listed at the end of this document.

**This file includes:**

Materials and Methods
Figs. S1 to S8
References (65–74)
Members of the Global Neurodegeneration Proteomics Consortium

### Materials and Methods

#### Study populations

**UK Biobank.** We analyzed 41,209 UK Biobank (UKB) participants with plasma proteomics measured at the baseline visit on the Olink Explore 3072 platform (29, 32). Participants were aged 40 to 70 at recruitment. After matching Olink identifiers to the compartment annotation, we removed proteins missing in more than 10% of samples and samples missing more than 10% of proteins, and we filled the remaining missing values with the mean of each protein computed across the whole cohort before the train and test split. Participants without a recorded visit age were excluded, leaving 41,209. A single random 80/20 partition (seed 42) assigned 32,967 participants to training and 8,242 to a held-out test set, and the same partition was used for every compartment.

**GNPC plasma.** The plasma arm of the Global Neurodegeneration Proteomics Consortium (GNPC) (30) included 19,051 participants with SomaScan v4.1 7K aptamer proteomics, aged 20 to 90. We trained organelle aging models independently in this cohort with the same approach as in UKB and tested associations with prevalent disease by logistic regression.

**GNPC CSF.** The GNPC cerebrospinal fluid (CSF) arm comprised 1,947 CSF samples measured with the same SomaScan v4.1 7K assay (30, 31), from participants aged 30 to 90. Organelle age estimates were available for 1,671 participants (one sample each), and all CSF analyses use these participants. We trained organelle aging models independently in this cohort and compared organelle aging between biofluids.

#### Ethics, consent and data governance

All analyses used previously collected, de-identified human cohort data obtained under the governance of the contributing studies; no new human samples were collected for this work. UK Biobank data were accessed under the application listed in the Acknowledgments. UK Biobank holds Research Tissue Bank approval from the North West Multi-centre Research Ethics Committee, and all participants gave written informed consent at recruitment. The GNPC data come from a federated release: each contributing cohort obtained ethics approval from its own institutional review board, written informed consent was obtained from participants or their legally authorized representatives, and all cohorts and data were de-identified by GNPC before release for analysis.

#### Organelle protein annotation

Proteins were assigned to subcellular compartments using the graph-based annotations of Hein et al. (28), who combined native organelle immunoprecipitation with mass spectrometry and k-nearest-neighbor graph clustering to localize more than 7,600 proteins. We merged the graph compartments into 13 modeled compartments, combining the nucleolus with the nucleus, the trans-Golgi with the Golgi, the early and recycling endosomes into one endosomal compartment and the actin cytoskeleton with the centrosome, and dropping the 14-3-3 scaffold, translation and P-body classes. Age models were fitted for all 13 compartments in UKB, and the 10 compartments with a held-out $R^2$ of at least 0.02 are carried through the paper. Ranked by held-out Pearson correlation with age, these are the plasma membrane (0.75), cytosol (0.57), cytoskeleton (0.51), Golgi (0.50), endosome (0.50), lysosome (0.49), ER (0.45), mitochondrion (0.25), nucleus (0.21) and ERGIC (0.21). The stress granule (0.10), peroxisome (0.08) and proteasome (0.06) fell below the threshold and appear only in GNPC panels, where they are marked. Some analyses use a subset of compartments, as stated in the figure legends. The proteins detectable on the Olink panel and passing quality control ranged from 4 (ERGIC) to 169 (cytosol) per compartment in UKB.

### Organelle age models

For each compartment in each cohort, we trained an elastic net regression model (65) to predict chronological age from z-scored abundances of the compartment's proteins. UKB models used $\alpha = 0.1$ and an L1 ratio of 0.5, were trained on the 32,967 training participants and were evaluated on the 8,242 held-out participants. In GNPC plasma, models were trained on 5,659 participants and evaluated on 1,415 held-out participants, with compartment-specific α between 0.010 and 0.100 and L1 ratios between 0.05 and 0.95. In GNPC CSF, α ($10^{-5}$ to $10^{1}$, seven log-spaced values) and the L1 ratio (0.01 to 0.99, nine values) were selected by grid search over five age-stratified cross-validation folds within the training set, scored by mean absolute error (MAE), and performance was evaluated on a held-out 20% test set by MAE, Pearson r and $R^2$.

CSF models were trained in participants free of cognitive impairment and neurological diagnosis (cognitive test score of at least 25 of 30, Clinical Dementia Rating of 0, and no recorded stroke, transient ischemic attack, traumatic brain injury, Alzheimer's disease, frontotemporal dementia, Parkinson's disease, amyotrophic lateral sclerosis, mild cognitive impairment or subjective cognitive impairment) and then applied to the whole cohort, so every CSF disease association compares cases against a model fitted without them. In UKB we applied no health-based exclusion, and the training set included prevalent cases.

The age gap is predicted minus chronological age. In GNPC CSF, it was taken relative to a locally weighted regression (LOWESS) (74) of predicted on chronological age fitted in cognitively healthy participants within each contributing cohort, and standardized to those participants. In GNPC plasma and UKB, the age gap was standardized within each compartment. For the UKB extreme-ager analyses, the age gap was additionally residualized on chronological age, as described below.

Second-generation models used the compartment age gaps as features in CoxNet survival analysis (scikit-survival CoxnetSurvivalAnalysis with an L1 penalty) to predict all-cause mortality in UKB. Hyperparameters were selected by threefold cross-validation over six L1 ratios (0.1 to 1.0) and 50 automatically generated α values. Performance was assessed by the concordance index, and each compartment received a mortality-risk score.

### Disease association analysis

**UKB (incident disease).** In the held-out test set, Cox proportional hazards models (66) related each compartment's predicted age (per SD) and mortality-risk score to time to incident disease, adjusted for chronological age and sex. Disease endpoints were defined from ICD-10 codes in linked hospital records. Twelve diseases, selected for sample size and clinical relevance, are shown in Fig. 2J; the full grid spans 49 endpoints and 13 compartments (637 tests), corrected by the Bonferroni method.

**GNPC (prevalent disease).** Using the GNPC models, we related organelle age gaps (per SD) to prevalent disease at the time of sampling by logistic regression adjusted for age and sex, separately in plasma and CSF. P values across the organelle-by-disease grid were adjusted by the Benjamini-Hochberg false discovery rate (FDR) (72).

**Biofluid comparison (Fig. 2, H and I).** For each biofluid, diagnosis and compartment we fitted one logistic model, diagnosis ~ age gap + age + sex, with the age gap restandardized within each analysis sample so that odds ratios are per SD in both biofluids. Controls were all participants without the diagnosis; participants with a missing diagnosis code, the plasma code 2 for diabetes and depression, or missing sex were excluded. P values were adjusted by the Benjamini-Hochberg method across the 52 tests within each biofluid. Plasma and CSF were compared for each diagnosis by a two-sided paired t-test of the absolute log odds ratio across the 13 compartments. No sample appears in both biofluid tables.

### Extreme ager analysis

For each compartment, the UKB extreme-ager score is the z-scored age gap residualized on chronological age by ordinary least squares in the training set and divided by its training-set standard deviation; the same coefficients and SD were applied to the held-out set. A compartment is accelerated when its score exceeds 2, decelerated when it falls below −2 and normal otherwise. Burden is the number of accelerated compartments among the ten carried forward. Survival analyses (Figs. 2C, 3C and 5D; fig. S3, A to C) use the held-out test set, exclude prevalent cases, cap follow-up at 15 years and estimate hazard ratios against the normal group, or against participants with no accelerated compartment for burden, by Cox regression with age and sex (lifelines, L2 penalty 0.01). Kaplan-Meier curves (71) show pointwise 95% confidence intervals.

Ager classes (Fig. 3, A and B) were assigned by rule: participants with exactly one accelerated compartment belong to that compartment's class, and participants with two or more belong to the multi-organelle class. The t-SNE map (perplexity 30, PCA initialization, random state 42) of the ten compartment scores is for display. In GNPC (Fig. 3D), burden counts compartments with a standardized age gap above 2 among nine compartments, and odds ratios come from logistic regression adjusted for age, age squared and sex. Disease-associated organelle signatures (Fig. 3E) were derived in GNPC plasma with a 90th-percentile threshold per compartment; for each disease, the three compartments with the highest disease specificity are shown, and diseases are ordered by Ward hierarchical clustering. Discordance (Fig. 3F) is the within-person SD across standardized organelle age gaps, modeled by logistic regression with the mean age gap, linear and quadratic age terms and sex.

### Functional module analysis

Principal component analysis was performed on the 41,209 × 13 matrix of age-detrended UKB compartment age gaps without rescaling the columns, and Fig. 5A plots the compartment loadings of the first two components. Their standard deviations range from 0.07 (proteasome) to 0.75 (plasma membrane), so both components reflect compartment variance as much as compartment function. Three functional modules were therefore assigned by compartment identity: synthesis (ER, ERGIC and Golgi), degradation (lysosome and endosome) and membrane (plasma membrane). The proteasome and the cytoskeleton enter no module. Module scores are the unweighted mean of the member compartments' extreme-ager scores and are not rescaled, so hazard ratios are per unit of module score. Cox models in the held-out test set included the three module scores jointly with age and sex, with follow-up capped at 15 years.

### Conditional models

To assess whether organelle-disease associations are independent of the other compartments, we fitted Cox models in the held-out test set for each compartment's age gap (per SD) with age and sex (unconditioned) and with the age gaps of the other 12 compartments added (conditioned). Unconditioned associations were corrected by the Bonferroni method across 91 tests; conditioned associations are shown at nominal $P < 0.05$.

### Mediation analysis

The contribution of lysosomal aging to the male excess in kidney disease (Fig. 5D) was assessed with the Baron-Kenny approach (67) in the held-out test set (8,092 participants, 738 incident kidney disease events). The total effect of male sex came from a Cox model with chronological age and sex (HR 1.58, 95% CI 1.36 to 1.83), and the direct effect from the same model with the lysosomal age gap added (HR 1.41, 95% CI 1.21 to 1.63). The proportion accounted for is the attenuation of the log hazard ratio, 1 − direct/total, which was 25% (95% CI 16 to 40%, 1,000 bootstrap resamples). This quantifies statistical accounting and does not establish causal mediation.

### Protein-disease driver analysis

For each disease-compartment pair, we estimated protein-level associations with disease by Cox models adjusted for age and sex in the full UKB cohort and plotted each protein's log hazard ratio against its weight in the compartment age model. Proteins with large positive values on both axes, ranked by the product of the absolute weight and $-\log_{10}(P)$, were considered drivers. Pathway enrichment used Enrichr (68) against Reactome (69) and KEGG (70), applied to the endosomal type 2 diabetes drivers (APP, BAG6, CLTA, IGF2R and NID1) and to the 16 ER kidney disease drivers.

### Protein interaction analysis

Within-compartment connectivity (Fig. 1C) was assessed against the STRING database (33). The human protein-links and protein-information tables of STRING release 12.0 (NCBI taxonomy 9606) were downloaded in bulk, edges were restricted to a combined score of at least 700, STRING identifiers were mapped to gene symbols, and the network was restricted to pairs in which both proteins carried a compartment annotation. Enrichment over random expectation was tested by permuting compartment labels. The ribbons in Fig. 4C are Pearson correlations between age-residualized protein abundances, not STRING edges.

### Aging velocity, protein drivers and pathway analysis

For each of the nine compartments with sufficient protein representation in GNPC plasma, member proteins were sign-aligned by the direction of their association with age, averaged into a single signature per participant and standardized across participants. Coordinated aging velocity is the absolute Pearson correlation between that signature and chronological age, divided by the SD of age and multiplied by ten, giving signature SDs per decade of age. Because the cohorts are cross-sectional, velocity is an age gradient across people rather than a rate of change within a person, and because alignment signs and velocities were estimated in the same participants, the ranking is descriptive. Intervals are the 2.5th and 97.5th percentiles over 1,000 bootstrap resamples of participants. The comparison in Fig. 4B is a one-sided subject bootstrap of the mean velocity of the lysosome, plasma membrane and Golgi minus the mean of the other six compartments.

Protein-level drivers within each compartment were ranked by the Pearson correlation of protein abundance with age, with significance at a Benjamini-Hochberg FDR below 0.05 across all measured proteins. Each imbalance index (Fig. 4, E, H and K) is the difference between the standardized abundances of two proteins, one rising and one falling with age, each among the most strongly age-associated proteins in its direction, chosen to represent the two arms of each compartment's remodeling: GM2A and ASAH1 (ganglioside clearance), SCARF2 and SRC (surface-receptor remodeling) and IGFBP2 and RAB6A (secretory trafficking). Indices were summarized within fixed age bins and compared between the youngest and oldest bins by two-sided Mann-Whitney U tests. Gene Ontology enrichment used Enrichr (68) against GO Biological Process 2021 for human, applied to the top-ranked age-associated proteins of each compartment; reported significance is the Enrichr adjusted P value.

### APOE genotype analyses

Associations with APOE ε4 dosage (0, 1 or 2 alleles) were estimated in genotyped GNPC CSF participants, 1,167 from contributor Q (figs. S4 and S6) and 1,378 pooled across contributors (fig. S5), by linear regression of standardized age gaps or protein abundances on dosage with age and sex and heteroskedasticity-consistent (HC3) standard errors. Contributions of individual proteins to a compartment's dosage coefficient were computed from the fixed model weights and the training standardization, so that they sum to the compartment association. Adjusted models added recorded AD, 36 immune and lipid proteins and four APOE assays. AD associations were estimated by logistic regression. P values were adjusted by the Benjamini-Hochberg method across the tested proteins, as stated in the legends.

### Statistical analysis

Analyses were performed in Python 3.12 with scikit-learn (elastic net), scikit-survival (penalized Cox models), lifelines (Cox models and Kaplan-Meier estimates), statsmodels, scipy and pandas, and figures were drawn with matplotlib. Nominal significance was set at $P < 0.05$, and multiplicity handling is stated for each analysis. Benjamini-Hochberg FDR control at $q < 0.05$ was applied to the GNPC organelle-by-disease grid (Fig. 2A, key * $q < 0.05$, ** $q < 0.01$, *** $q < 0.001$), the biofluid odds ratios (Fig. 2H), the extreme-tertile and cognitive-impairment odds ratios (Fig. 2, B and F), the APOE dosage regressions and the protein-level age associations (Fig. 4, D, G and J). Bonferroni correction was applied to the UKB organelle-by-disease grid (637 tests) and to the unconditioned associations of Fig. 5C (91 tests). The within-AD severity contrast (Fig. 2D, two-sided Mann-Whitney U tests across ten compartments) and the disease-signature odds ratios (Fig. 3E) carry no correction. UKB extreme-ager, burden, module, conditional and mediation analyses use the held-out test set, whereas the protein-level driver models use the full cohort. Protein-wise mean imputation preceded the train and test split, and the mortality models take the age gaps as their only inputs, so age and mortality analyses are not independent evidence.

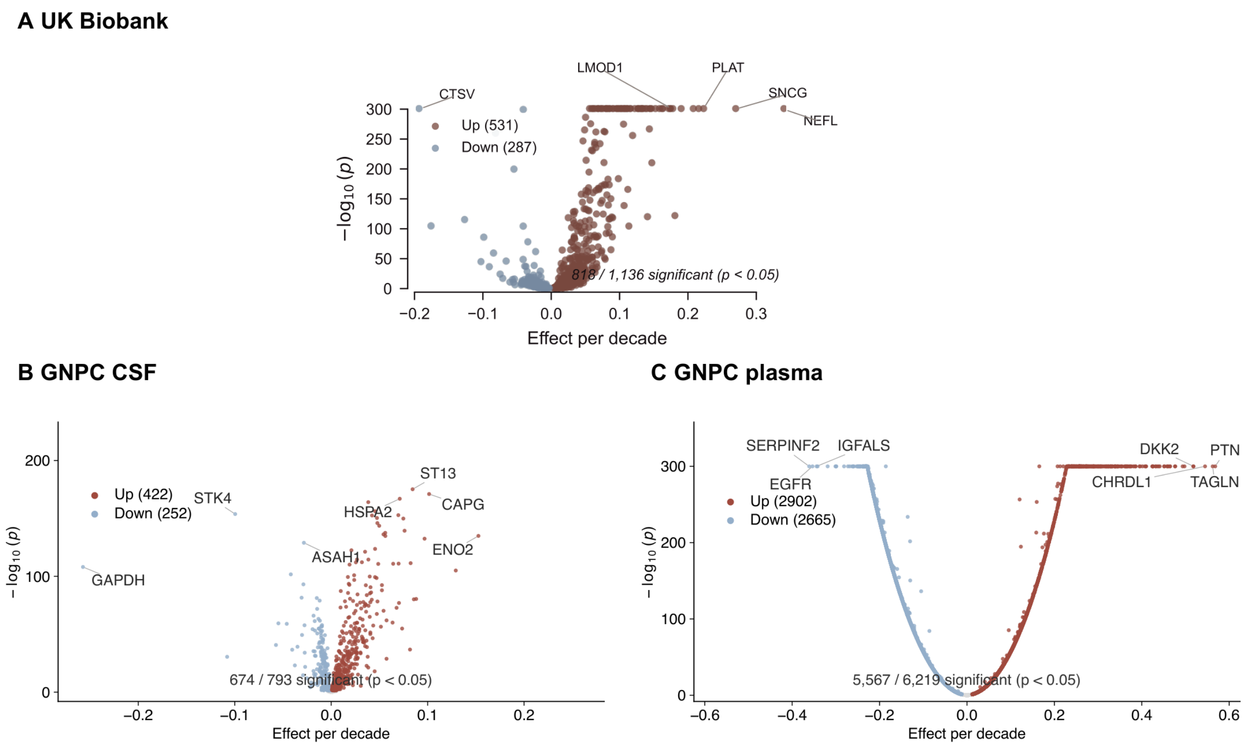


**Fig. S1. Protein associations with chronological age across cohorts.** (**A** to **C**) Protein-level age associations in UKB (A), GNPC CSF (B) and GNPC plasma (C). Points represent assayed proteins; the horizontal coordinate is the regression slope per decade of chronological age and the vertical coordinate is −log10(P). Red and blue indicate nominally significant positive and negative associations, respectively (two-sided $P < 0.05$); gray denotes other proteins. Counts are 531 positive and 287 negative among 1,136 tested proteins in A, 422 and 252 among 793 in B, and 2,902 and 2,665 among 6,219 in C. Panels B and C use separate regressions of protein abundance on chronological age.

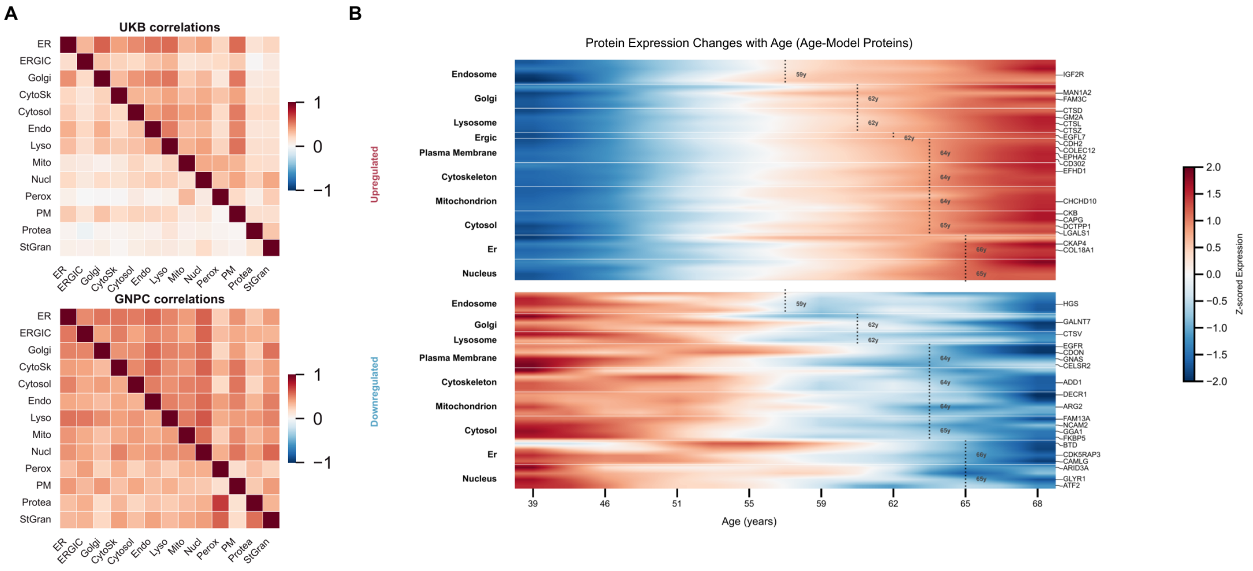


**Fig. S2. Correlations between organelle age gaps and age-associated changes in age-model proteins.** (**A**) Pairwise correlations between organelle age gaps in UKB (upper matrix) and GNPC plasma (lower matrix). Color indicates the correlation coefficient, with red denoting positive and blue negative correlations. Compartments are arranged in the same order along both axes. (**B**) Age-binned, standardized abundance of proteins selected by the chronological-age models, grouped by organelle. Upper and lower heatmaps display proteins increasing and decreasing with age, respectively. Rows represent proteins, columns represent chronological-age bins, and color indicates standardized abundance. Selected protein names are shown at right; dotted lines mark the annotated ages within each compartment. ER, endoplasmic reticulum; ERGIC, ER–Golgi intermediate compartment; PM, plasma membrane.

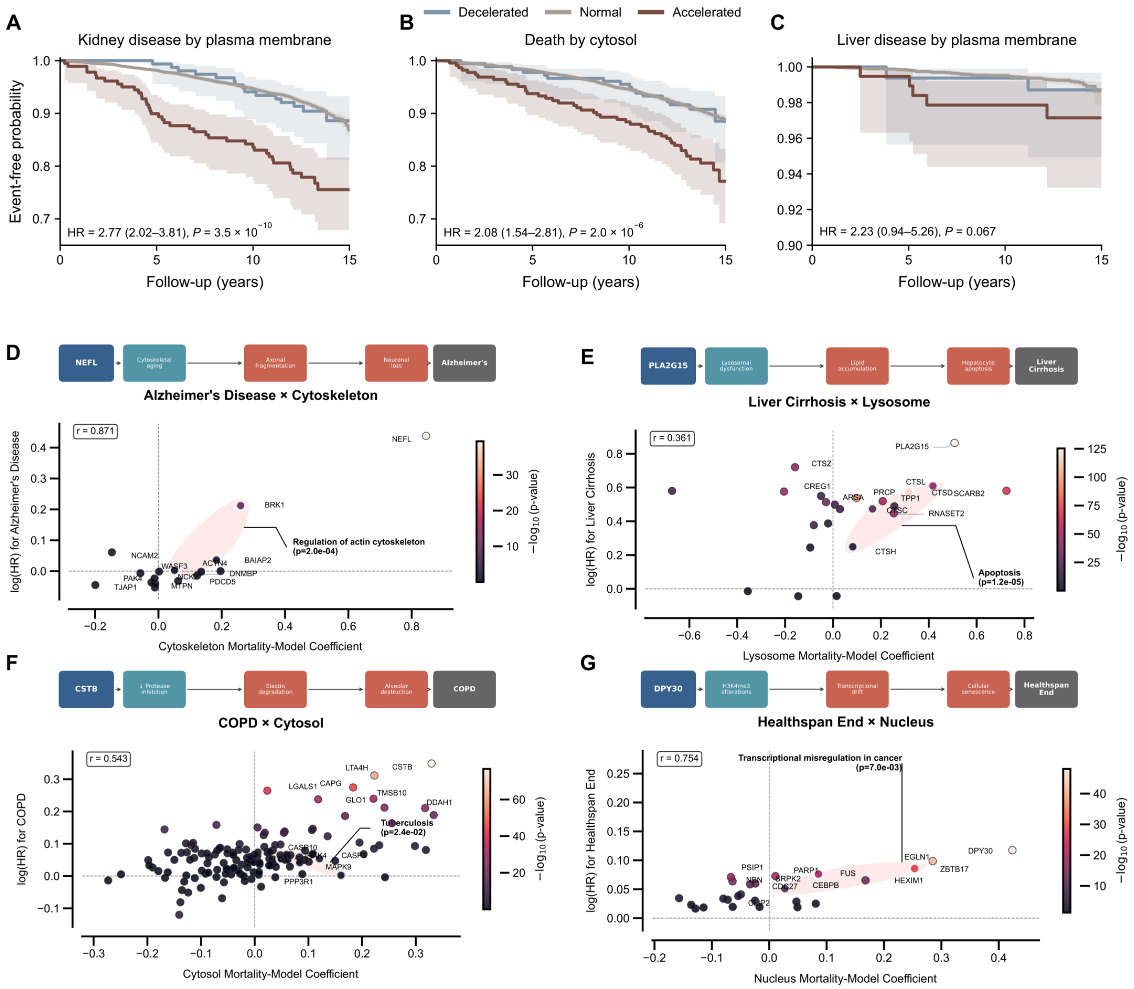


**Fig. S3. Held-out survival and protein associations for selected disease–organelle pairs.** (**A** to **C**) Kidney disease and plasma membrane (A), death and cytosol (B) and liver disease and plasma membrane (C) in the UKB held-out test set (8,242 participants before endpoint exclusions). Groups use the extreme-ager score of Fig. 2C: accelerated, score > 2; normal, −2 to 2; decelerated, < −2. Lines show Kaplan–Meier estimates (71); shading denotes pointwise 95% CIs. Follow-up is capped at 15 years and prevalent cases are excluded. Annotations give age- and sex-adjusted accelerated-versus-normal Cox HRs, 95% Wald CIs and two-sided Wald P values (L2 penalty 0.01). Events/participants are 40/185, 44/224 and 5/197 in the accelerated groups, 683/7,749, 756/7,839 and 64/7,852 in the normal groups and 15/158, 17/179 and <5/161 in the decelerated groups (A, B and C, respectively). (**D** to **G**) Protein-level disease log(HR) versus mortality-model coefficient for AD and cytoskeleton (D), liver cirrhosis and lysosome (E), chronic obstructive pulmonary disease and cytosol (F) and healthspan end and nucleus (G). Color denotes association −log10(P); selected proteins and enriched pathways are labeled. Pearson r summarizes the displayed coefficient–association relationship. Schematics above each plot depict proposed links between proteins, pathways and disease.

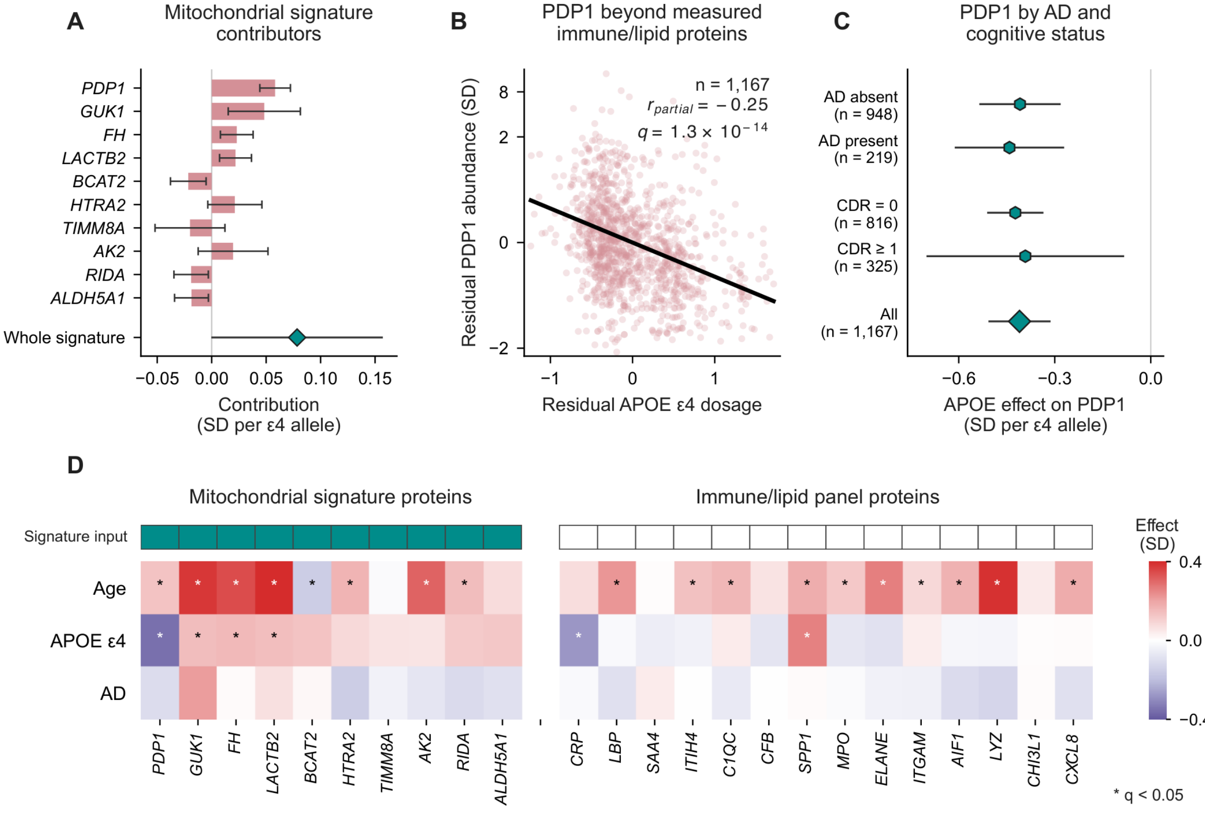


**Fig. S4. APOE ε4-associated PDP1 variation within the mitochondrial signature.** (**A**) Ten largest absolute protein contributions to the AD-adjusted ε4 dosage coefficient of the mitochondrial signature in 1,167 genotyped CSF participants from contributor Q. Bars show signed contributions and 95% heteroskedasticity-consistent (HC3) intervals; the diamond denotes the complete 82-input signature. Contributions use fixed weights and training standardization. (**B**) Partial association between ε4 dosage and PDP1 after conditioning on age, sex, recorded AD, 36 immune/lipid proteins and four APOE assays. Both variables are residualized against identical covariates. The vertical axis is symmetric-log, linear within ±1 SD. The coefficient is −0.410 SD per allele (95% CI −0.507 to −0.313); q is Benjamini–Hochberg-adjusted across 110 mitochondrial and lysosomal inputs (72). (**C**) PDP1 dosage coefficients and 95% HC3 intervals within AD and CDR strata and overall. AD-stratified models include age, sex and 40 protein covariates; CDR-stratified and overall models additionally include AD status. Dosage-by-AD interaction P = 0.915. (**D**) Joint age, dosage and AD coefficients for the same ten mitochondrial inputs and 14 immune proteins. Outcomes are standardized; age effects are per decade and dosage effects per ε4 allele. Asterisks indicate q < 0.05 across 146 proteins separately for each contrast.

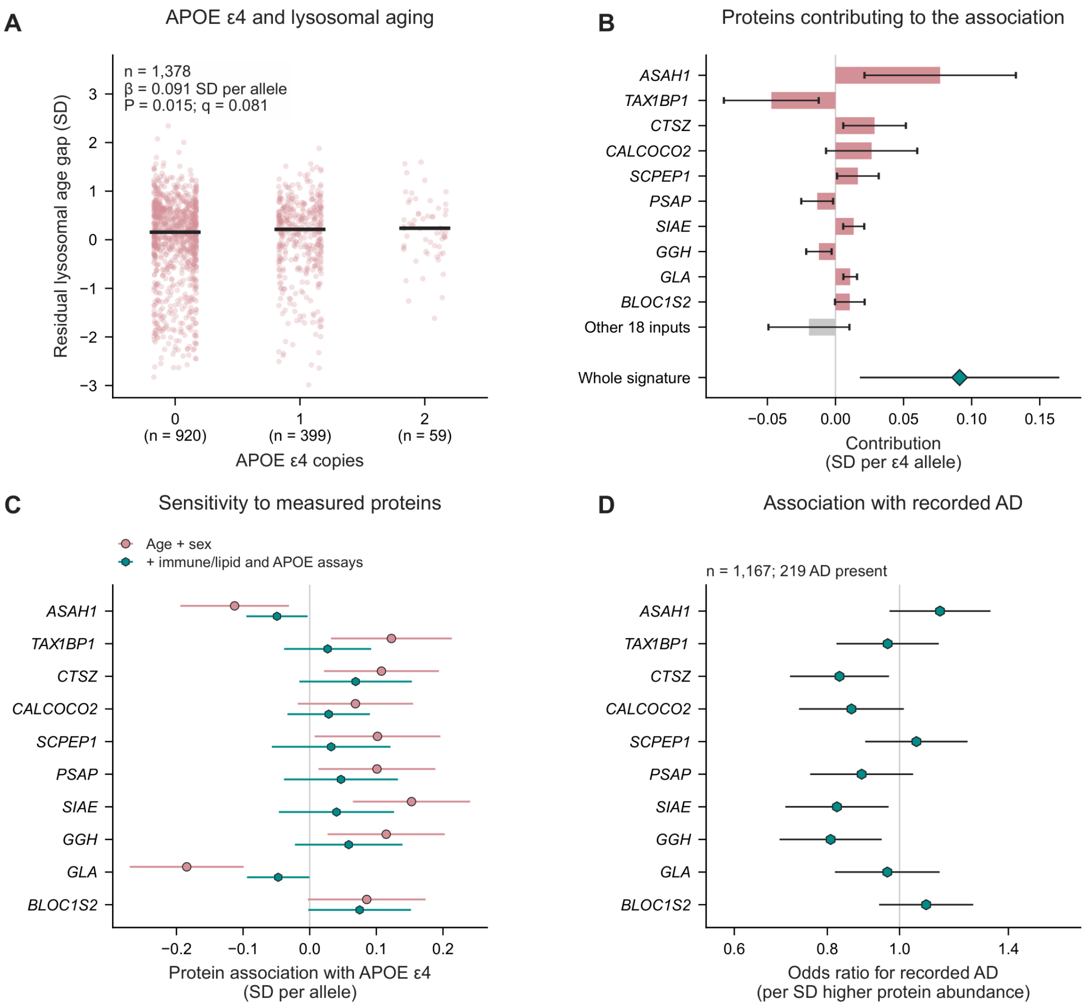


**Fig. S5. Pooled APOE ε4 associations with the lysosomal signature and its inputs.** (**A**) Age- and sex-residualized lysosomal age gaps by ε4 dosage in 1,378 genotyped participants (0 copies, n = 920; 1 copy, n = 399; 2 copies, n = 59). Observations are shown with horizontal jitter and group medians. The age- and sex-adjusted dosage coefficient is 0.091 SD per allele (95% HC3 CI 0.018–0.164; P = 0.0147; Benjamini–Hochberg q = 0.0809 across 11 compartments). (**B**) Ten largest absolute signed protein contributions, the combined contribution of the remaining 18 inputs, and the complete-signature coefficient. All 28 contributions sum to the whole-signature association; 25 fitted weights are nonzero. (**C**) Dosage coefficients for the same ten proteins, adjusted for age and sex alone or additionally for 36 immune/lipid proteins and four APOE assays. Lines are 95% HC3 intervals. (**D**) Recorded AD associations for these proteins in 1,167 participants from contributor Q, including 219 with AD. Points and lines denote logistic ORs per SD and 95% Wald CIs adjusted for age, sex and ε4 dosage.

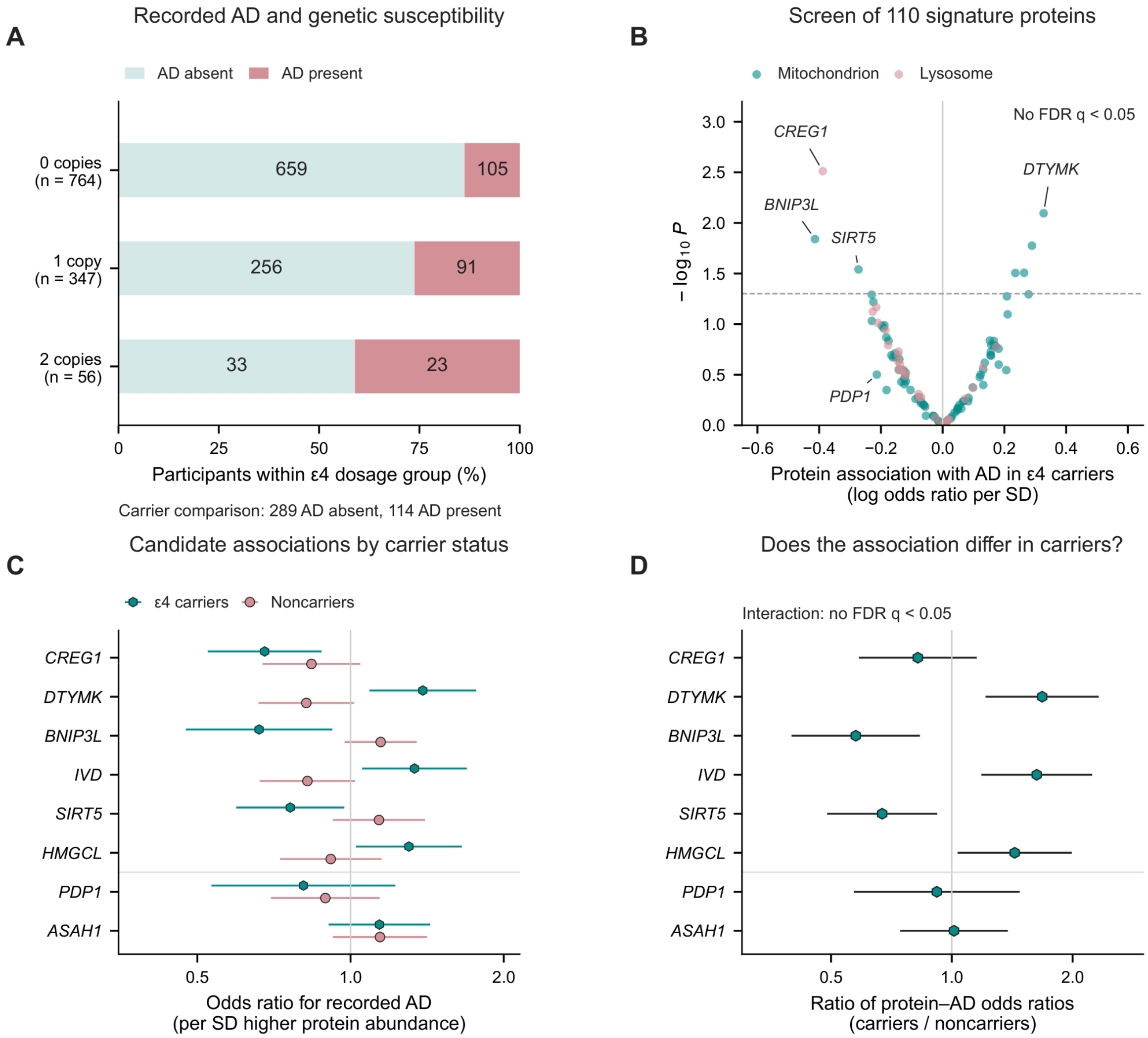


**Fig. S6. Protein associations with recorded AD among APOE ε4 carriers.** APOE ε4 dosage is an established AD susceptibility factor (73). (**A**) Recorded AD proportions and counts by ε4 dosage in 1,167 genotyped participants: 403 carriers (114 AD present, 289 absent) and 764 noncarriers (105 present, 659 absent). (**B**) Logistic association screen of 110 mitochondrial and lysosomal signature inputs among carriers, adjusted for age, sex and ε4 homozygosity. Coordinates are log(OR) per SD and −log10(two-sided Wald P); the dashed line marks nominal P = 0.05. No protein passed Benjamini–Hochberg q < 0.05. (**C**) Carrier and noncarrier associations for the six proteins with the smallest carrier P values, plus PDP1 and ASAH1 as prior candidates irrespective of significance. Abundance is standardized in the full sample; points and lines show ORs and pointwise 95% Wald intervals. (**D**) Protein-by-carrier interaction OR ratios and 95% Wald intervals from joint logistic models including age, sex, carrier status, homozygosity, protein abundance and the interaction. Interaction P values use a one-degree-of-freedom likelihood-ratio test, with Benjamini–Hochberg correction across 110 tests.

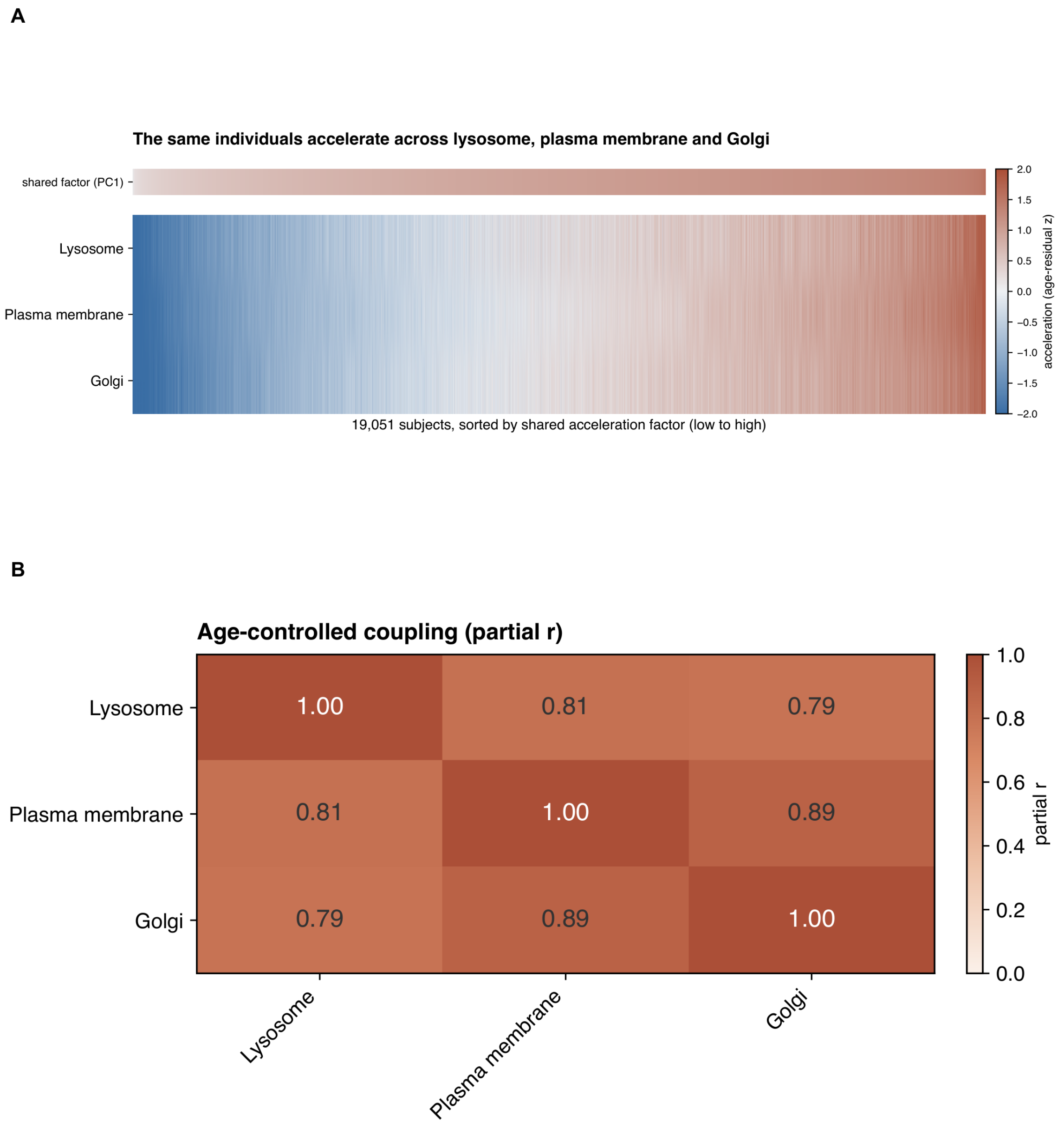


**Fig. S7. Shared age-residualized lysosomal, plasma-membrane and Golgi signatures.** (**A**) Standardized residual protein signatures for lysosome, plasma membrane and Golgi in 19,051 GNPC plasma participants, sorted by their first principal-component score. Protein z scores were aligned to their age-association direction and averaged within each compartment; locally weighted scatterplot smoothing (LOWESS) age trends (fraction 0.4) were subtracted and residuals standardized (74). The top strip shows the shared component, oriented toward positive mean loadings. Colors in the compartment matrix are displayed between −2 and +2 residual SD. (**B**) Pearson correlations of the age-detrended signatures.

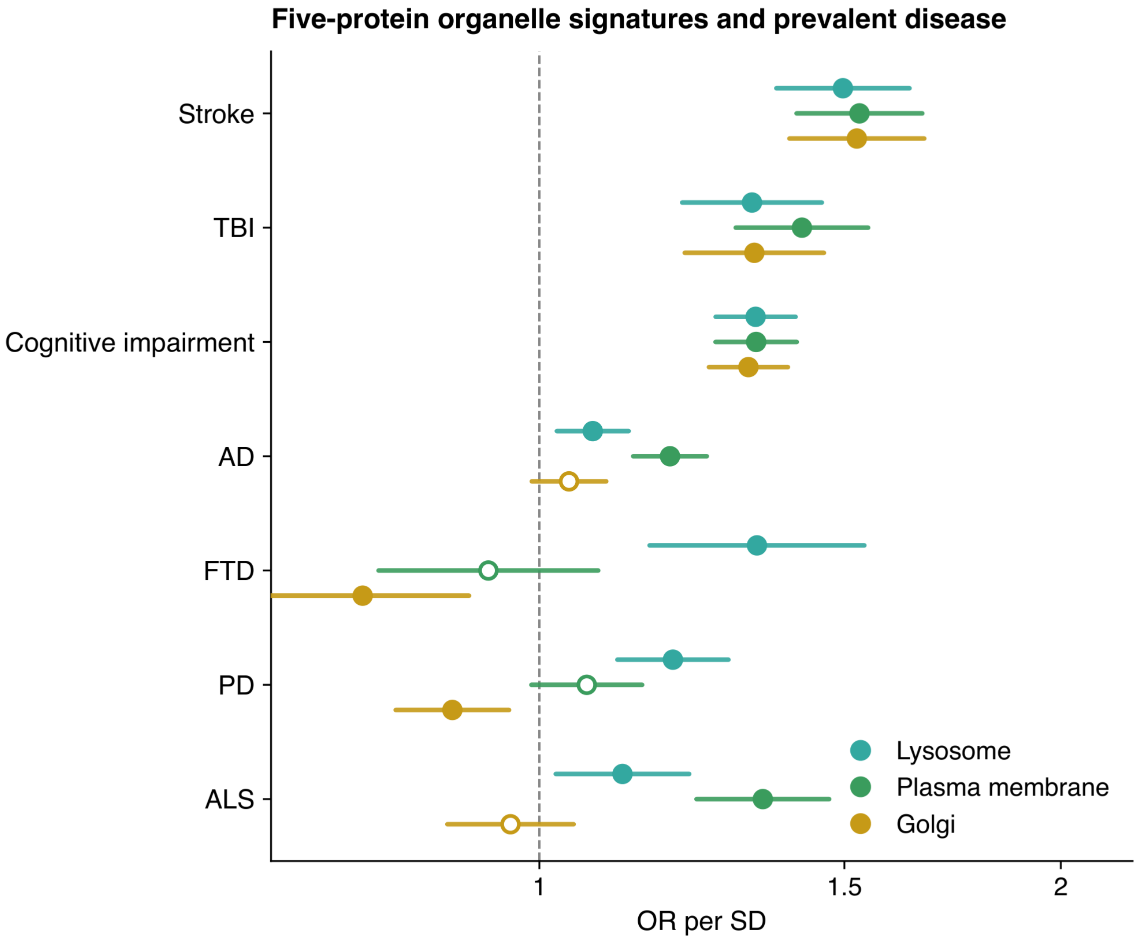


**Fig. S8. Five-protein organelle signatures and prevalent neurological disease.** ORs per SD of lysosomal, plasma-membrane and Golgi signatures for seven diagnoses in GNPC plasma. Each signature comprises the five proteins with the largest absolute age correlations within that compartment, aligned to the direction of aging, averaged, linearly age-detrended and standardized. Separate logistic models include age and categorical sex. The reference group has CDR = 0 and none of the specified AD, Parkinson's disease (PD), amyotrophic lateral sclerosis (ALS), frontotemporal dementia (FTD), cognitive impairment, stroke, transient ischemic attack or traumatic brain injury (TBI) diagnoses. Points and lines denote ORs and 95% Wald CIs; filled points indicate nominal $P < 0.05$ and open points $P \geq 0.05$.

### Members of the Global Neurodegeneration Proteomics Consortium (GNPC)


Charles H Adler, Mayo Clinic Arizona, Scottsdale, Arizona, USA

Alireza Atri, Banner Sun Health Research Institute, Sun City, Arizona, USA

Thomas G Beach, Banner Sun Health Research Institute, Sun City, Arizona, USA

Graham Bearden, Alzheimer's Disease Data Initiative, Kirkland, WA

James D. Berry, Sean M. Healey and AMG Center for ALS, Neurology

Merce Boada, Ace Alzheimer Center Barcelona, Universitat Internacional de Catalunya, 08029 Barcelona, Spain; Biomedical Research Networking Centre in Neurodegenerative Diseases (CIBERNED), National Institute of Health Carlos III, 28029 Madrid, Spain

Merle Bode, Hertie Institute for Clinical Brain Research, Neurodegenerative Diseases, Tübingen; German Center of Neurodegenerative Diseases, Department of Neurodegenerative Diseases, Tübingen

Bradley Boeve, Mayo Clinic, Neurology Department, Rochester, MN

Hillary Bounds, Gates Ventures, Seattle, WA

Alfredo Cabrera-Socorro, Johnson & Johnson, NS TA, Beerse, Belgium

Amanda Fernandez Cano, Ace Alzheimer Center Barcelona, Universitat Internacional de Catalunya, 08029 Barcelona, Spain; Biomedical Research Networking Centre in Neurodegenerative Diseases (CIBERNED), National Institute of Health Carlos III, 28029 Madrid, Spain

Kaitlin B. Casaletto, University of California, San Francisco, Neurology Department, San Francisco, CA

Richard J Caselli, Mayo Clinic Arizona, Scottsdale, Arizona, USA

Yike Chen, Washington University School of Medicine, Department of Psychiatry, St. Louis, 63110, MO, USA.; NeuroGenomics and Informatics Center, Washington University School of Medicine, St. Louis, 63110, MO, USA.

Matthew H.S. Clement, Alzheimer's Disease Data Initiative, Kirkland, WA

Eric B. Dammer, Emory University School of Medicine, Atlanta, GA, USA; Emory University School of Medicine, Department of Biochemistry, Atlanta, GA, USA

Sterre de Boer, Alzheimer Center Amsterdam, Neurology, Amsterdam UMC, Amsterdam, the Netherlands; Amsterdam Neuroscience, Amsterdam, the Netherlands

Niels De Meirleir, Johnson & Johnson, NS TA, Beerse, Belgium

Marta del Campo Milan, Barcelonaβeta Brain Research Center (BBRC), Pasqual Maragall Foundation, Barcelona, Spain; Hospital del Mar Research Institute, Barcelona, Spain

Duc Duong, Emory University School of Medicine, Atlanta, GA, USA; Emory University School of Medicine, Department of Biochemistry, Atlanta, GA, USA

Maria Victoria Fernandez, Ace Alzheimer Center Barcelona, Universitat Internacional de Catalunya, 08029 Barcelona, Spain

Lawrence Fourgeaud, Johnson & Johnson, NS TA, La Jolla, USA

Raquel Puerta Fuentes, Ace Alzheimer Center Barcelona, Universitat Internacional de Catalunya, 08029 Barcelona, Spain; PhD Program in Biotecnology, Faculty of Pharmacy and Food Sciences, University of Barcelona, 08028 Barcelona, Spain

Jordan Fuller, Gates Ventures, Seattle, WA

Su Gao, Indiana Alzheimer's Disease Research Center, Indianapolis, IN; Indiana University School of Medicine, Department of Biostatistics & Health Data Science, Indianapolis, IN

John Gibbons, Rush Alzheimer's Disease Center, Department of Neurological Sciences, Chicago, IL, USA

Pablo Garcia Gonzalez, Ace Alzheimer Center Barcelona, Universitat Internacional de Catalunya, 08029 Barcelona, Spain; Biomedical Research Networking Centre in Neurodegenerative Diseases (CIBERNED), National Institute of Health Carlos III, 28029 Madrid, Spain

Gyujin Heo, Washington University School of Medicine, Department of Psychiatry, St. Louis, 63110, MO, USA.; NeuroGenomics and Informatics Center, Washington University School of Medicine, St. Louis, 63110, MO, USA.

Hilary Heuer, University of California, San Francisco, Neurology Department, San Francisco, CA

Liping Hou, Johnson & Johnson, Spring House, USA

Yen-Ning Huang, Indiana Alzheimer's Disease Research Center, Indianapolis, IN; Indiana University School of Medicine, Department of Radiology & Imaging Sciences, Indianapolis, IN

Clifford R. Jack, Jr, Mayo Clinic, Radiology

Emily Kogan, Johnson & Johnson, JRD DSDH, Cambridge, USA

Jessica B Langbaum, Banner Alzheimer's Institute, Phoenix, Arizona, USA

Argentina Lario-Lago, University of California, San Francisco, Neurology Department, San Francisco, CA

Shuwei Li, Johnson & Johnson, Spring House, USA

Shiwei Liu, Indiana Alzheimer's Disease Research Center, Indianapolis, IN; Indiana University School of Medicine, Department of Radiology & Imaging Sciences, Indianapolis, IN

Menghan Liu, Washington University School of Medicine, Department of Psychiatry, St. Louis, 63110, MO, USA.; NeuroGenomics and Informatics Center, Washington University School of Medicine, St. Louis, 63110, MO, USA.

Marta Marquie, Ace Alzheimer Center Barcelona, Universitat Internacional de Catalunya, 08029 Barcelona, Spain; Biomedical Research Networking Centre in Neurodegenerative Diseases (CIBERNED), National Institute of Health Carlos III, 28029 Madrid, Spain

Caitlin P. McHugh, Alzheimer's Disease Data Initiative, Kirkland, WA

Martine Meyer, Johnson & Johnson, NS TA

Silke Miller, Johnson & Johnson, NS TA, La Jolla, USA

Elizabeth Mlynarski, Johnson & Johnson, JRD DSDH, Spring House, USA

Diederik Moechars, Johnson & Johnson, NS TA, Beerse, Belgium

Patricia Moran-Losada, Stanford University, The Phil and Penny Knight Initiative for Brain Resilience, Stanford, CA, USA; Stanford University, Wu Tsai Neurosciences Institute, Stanford, CA, USA; Stanford University School of Medicine, Department of Neurology and Neurological Sciences, Stanford, CA, USA

Paige Opsahl, Gates Ventures, Seattle, WA

Tamina Park, Indiana Alzheimer's Disease Research Center, Indianapolis, IN; Indiana University School of Medicine, Department of Radiology & Imaging Sciences, Indianapolis, IN

Mukta Phatak, Alzheimer's Disease Data Initiative, Kirkland, WA

Joni Lindbohm, MD, PhD, University College London, UCL Brain Sciences, London, UK; University of Helsinki, Clinicum, Helsinki, Finland

Joseph Pick, Johnson & Johnson, Spring House, USA

Yolande AL Pijnenburg, Alzheimer Center Amsterdam, Neurology Department, Amsterdam, the Netherlands; Amsterdam Neuroscience, Amsterdam, the Netherlands

Michael Price, Michael J. Fox Foundation, New York, NY, USA

Shannon Risacher, Indiana Alzheimer's Disease Research Center, Indianapolis, IN; Indiana University School of Medicine, Department of Radiology & Imaging Sciences, Indianapolis, IN

Julio C. Rojas, University of California, San Francisco, Neurology Department, San Francisco, CA

Howard J. Rosen, University of California, San Francisco, Neurology Department, San Francisco, CA

Tamsin Sargood, Johnson & Johnson, Global Development, UK

Claudia Schulte, Hertie Institute for Clinical Brain Research, Neurodegenerative Diseases, Tübingen; German Center of Neurodegenerative Diseases, Department of Neurodegenerative Diseases,

Tübingen

Weiwei Schultz, Johnson & Johnson, JRD DSDH, Titusville, USA

Geidy E Serrano, Banner Sun Health Research Institute, Sun City, Arizona, USA

Nicholas T. Seyfried, Emory University School of Medicine, Atlanta, GA, USA; Emory University School of Medicine, Department of Neurology, Atlanta, GA, USA; Emory University School of Medicine, Department of Biochemistry, Atlanta, GA, USA

Todd Sherer, Michael J. Fox Foundation, New York, NY, USA

Emily Smith, Indiana Alzheimer's Disease Research Center, Indianapolis, IN; Indiana University School of Medicine, Department of Radiology & Imaging Sciences, Indianapolis, IN

Adam M. Staffaroni, University of California, San Francisco, Neurology Department, San Francisco, CA

Russell H. Swerdlow, University of Kansas Alzheimer's Disease Research Center, Kansas City, Kansas, USA; University of Kansas, Neurology, Kansas City, Kansas, USA

Shinya Tasaki, Rush Alzheimer's Disease Center, Department of Neurological Sciences, Chicago, IL, USA

Charlotte Teunissen, Neurochemistry Laboratory, Neurology Department, Amsterdam, the Netherlands; Amsterdam Neuroscience, Amsterdam, the Netherlands

Terri G. Thompson, OnPoint Scientific, Inc, San Diego, CA, USA

Qu Tian, NIH/NIA

Jigyasha Timsina, Washington University School of Medicine, Department of Psychiatry, St. Louis, 63110, MO, USA.; NeuroGenomics and Informatics Center, Washington University School of Medicine, St. Louis, 63110, MO, USA.

Abolfazl Doostparast torshizi, Johnson & Johnson, Spring House, USA

Sergi Valero, Ace Alzheimer Center Barcelona, Universitat Internacional de Catalunya, 08029 Barcelona, Spain; Biomedical Research Networking Centre in Neurodegenerative Diseases (CIBERNED), National Institute of Health Carlos III, 28029 Madrid, Spain

Wiesje M van der Flier, Alzheimer Center Amsterdam, Neurology Department, Amsterdam, the Netherlands; Amsterdam Neuroscience, Amsterdam, the Netherlands; Epidemiology and Data Science, Amsterdam UMC

Julia D. Webb, University of California, San Francisco, Neurology Department, San Francisco, CA

Bryan K Woodruff, Mayo Clinic Arizona, Scottsdale, Arizona, USA

Ying Xu, Washington University School of Medicine, Department of Psychiatry, St. Louis, 63110, MO, USA.; NeuroGenomics and Informatics Center, Washington University School of Medicine, St. Louis, 63110, MO, USA.

Mariet A. Younkin, Mayo Clinic, Neurology, Rochester, MN
